# Assessing the potential of BirdNET to infer European bird communities from large-scale ecoacoustic data

**DOI:** 10.1101/2023.12.06.570351

**Authors:** David Funosas, Luc Barbaro, Laura Schillé, Arnaud Elger, Bastien Castagneyrol, Maxime Cauchoix

## Abstract

1. Passive acoustic monitoring has become increasingly popular as a practical and cost-effective way of obtaining highly reliable acoustic data in ecological research projects. Increased ease of collecting these data means that, currently, the main bottleneck in ecoacoustic monitoring projects is often the time required for the manual analysis of passively collected recordings. In this study we evaluate the potential and current limitations of BirdNET-Analyzer v2.4, the most advanced and generic deep learning algorithm for bird recognition to date, as a tool to assess bird community composition through the automated analysis of large-scale ecoacoustic data.
2. To this end, we study 3 acoustic datasets comprising a total of 629 environmental soundscapes collected in 194 different sites spread across a 19° latitude span in Europe. We analyze these recordings both with BirdNET and by manual listening by local expert birders, and we then compare the results obtained through the two methods to evaluate the performance of the algorithm both at the level of each single vocalization and for entire recording sequences (1, 5 or 10 min).
3. Our analyses reveal that BirdNET identifications can be highly reliable if a sufficiently high minimum confidence threshold is used. However, the current recall of the algorithm is markedly low when the minimum confidence threshold is adjusted to ensure high levels of precision. Thus, we found that F1-scores remain moderate (<0.5) for all datasets and confidence thresholds studied. We therefore estimate that acoustic datasets of extended duration are currently necessary for BirdNET to provide a reliable and minimally comprehensive picture of the target bird community. Our results also suggest that BirdNET performance is not significantly influenced by the type of recorder used or the habitat recorded but is modulated by the volume of species-specific acoustic data available online.
4. We conclude that a judicious use of AI-based IDs provided by BirdNET can represent a novel and powerful method to assist in the assessment of bird community composition through the automated analysis of large-scale ecoacoustic data. Finally, we provide best use recommendations to ensure optimal results from the algorithm for the ecological study of bird communities.

## 1. INTRODUCTION

In recent years, passive acoustic monitoring (PAM) based on autonomous recording units (ARUs) has become an increasingly popular option for monitoring avian populations and communities (Shonfield & Bayne, 2017). Compared to active acoustic monitoring (AAM) methods such as point counts or active searches, ARUs could provide a more cost-effective way of obtaining data (Melo et al., 2021; Darras et al., 2018) and a higher degree of standardization in the data collection process (Darras et al. 2019). Moreover, ARUs offer the possibility of listening to distant, noisy or dubious animal sounds multiple times, potentially facilitating their identification (Sugai et al., 2019). At the same time, ARUs make it possible to collect substantial amounts of ecoacoustic data in a non-invasive way for local wildlife.

ARUs can be especially useful for biodiversity monitoring projects in hard-to-reach areas, where difficult access conditions might hinder frequent data collection through AAM methods, and might improve the detectability of species whose vocal activity patterns do not coincide with the dates and times at which active monitoring is usually conducted (Sebastián-González et al., 2018). This greater facility to collect ecoacoustic data at regular time intervals in a standardized way makes it possible to study the variations in vocal activity patterns of one or more species along a given time gradient (e.g., time of day, week of the year) (Gasc et al., 2013; Towsey et al., 2014). Additionally, the analysis of soundscape variation across time and space allows to study potential correlations between the presence of anthropogenic sounds and the acoustic activity of sound-producing animals (Blumstein et al., 2011). This means that PAM could serve as a reliable way to obtain complementary information to cover the gaps left by traditional monitoring methods (Bradfer-Lawrence et al., 2023). Indeed, recent studies suggest that the combined use of PAM and AAM may provide a more accurate picture of the presence and abundance of target species than the use of only one of the two methods (Bobay et al., 2018; Shaw et al., 2021).

The increased ease of obtaining large amounts of ecoacoustic data through PAM means that, currently, the main bottleneck in ecoacoustic monitoring is often the time required for the analysis of passively collected data (Symes et al., 2022). Manual analysis of recordings by experts requires time-intensive dedication and therefore severely limits the amount of data that can be processed. Thus, the automated analysis of recordings could represent a potential solution to the current disparity between the efforts required to obtain data through PAM and the efforts necessary to manually analyze these data. In recent years, deep learning techniques have allowed for significant advances in the development of different tools for the automated recognition of biological sounds (Stowell, 2021). However, this field is still in a developing phase, and the use of these algorithms often results in significant numbers of false positives or false negatives, thus requiring subsequent manual validation (Knight et al., 2017). Hence, the development of ecoacoustic data processing algorithms that can reliably identify the species recorded without manual supervision could allow for a substantial transformation of the methods through which biodiversity monitoring projects are conducted (Liu et al., 2022).

In this study we evaluate BirdNET-Analyzer v2.4, a deep neural network algorithm developed by the Cornell Lab of Ornithology which is capable of identifying 6512 bird species worldwide based on their sounds. BirdNET, in addition to detecting and identifying bird sounds present in a given recording, assigns a confidence score between 0 and 1 to each identification (Kahl et al., 2021), thus providing information about the reliability of its results. If considered sufficiently reliable, BirdNET could partially or even fully automate the analysis of ample amounts of ecoacoustic data that can already be efficiently obtained with ARUs (Stowell, 2021; Borowiec et al., 2022). Moreover, the recently released BirdNET App (Wood et al., 2022) might further contribute to the collection of ornithological data by helping non-experts identify bird vocalizations heard on the field.

Several recent studies have tested the potential and limitations of BirdNET as a tool to assess bird community composition through the automated analysis of environmental soundscapes (Kahl et al., 2021; Arif et al., 2020; Brüggemann et al., 2021; Tolkova et al., 2021; Sethi et al., 2021; Toenies & Rich, 2021; Cole et al., 2022; Höchst et al., 2022; Malamut, 2022). However, as highlighted in a recent review of these previous studies (Pérez-Granados, 2023), multiple analyses of potential relevance are still lacking. More concretely, the impact of (i) the number of species vocalizing simultaneously, (ii) the recording environment (i.e., the soundscape or acoustic habitat type), (iii) the type of recorder employed, (iv) the volume of species-specific acoustic data available online, and (v) the input values for the overlap and detection sensitivity parameters of the algorithm on BirdNET performance remains to be evaluated. Moreover, the review suggests that BirdNET performance should be studied not only at the entire recording level but also at the level of each single identification/vocalization. Last but not least, BirdNET detection capacity (i.e., recall rate) across a large range of European soundscapes has yet to be investigated.

To fill this knowledge gap we analyze, both automatically with BirdNET and manually by expert birders, 629 environmental soundscapes collected in 194 different sites spread across a 19° latitude gradient in Europe (Fig. 1). We then compare the identifications obtained with BirdNET against those made by experts in order to assess the performance of the algorithm in terms of precision and recall (see subsection 2.4.1). Subsequently, we examine how to optimize the existing trade-off between these two metrics and how different features of the acoustic dataset analyzed might influence BirdNET performance. Finally, we provide best use recommendations to ensure optimal results from the algorithm for the ecological study of bird communities.

**Fig. 1:**
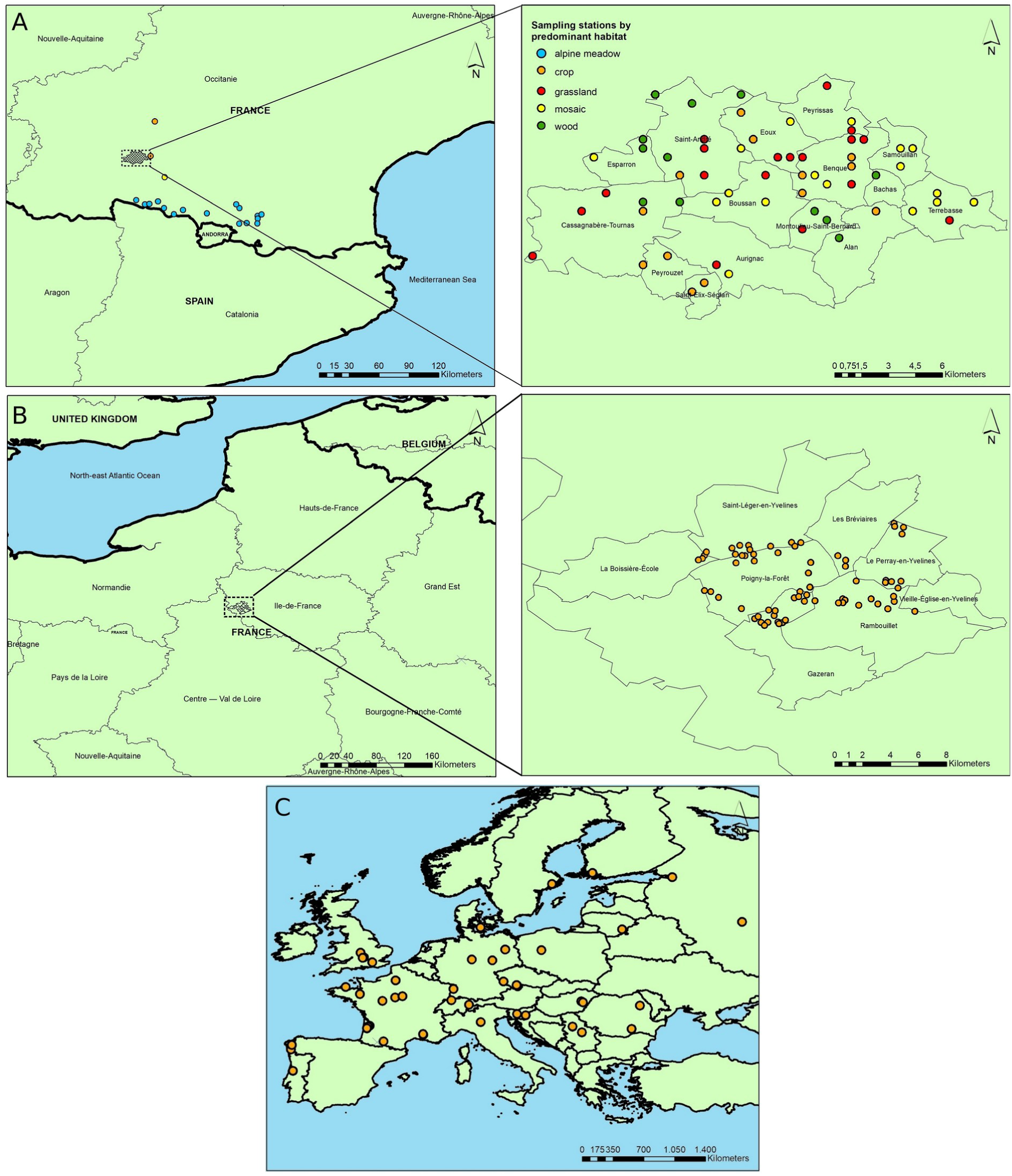
Maps showing (a) the 79 sampling stations in the ZA-Pygar dataset, (b) the 68 sampling stations in the Rambouillet dataset, and (c) the 47 sampling stations in the TreeBodyguards dataset. The predominant habitat in each station is only specified for the ZA-Pygar dataset.

## 2. METHODS

### 2.1. Study area and soundscape collection

The environmental soundscapes analyzed in this study originated from three independent acoustic datasets (Table 1, Appendix S1). Our first dataset, hereafter referred to as ZA-Pygar, consists of a total of 237 1-minute recordings collected from 2019 to 2022 in 79 different sampling sites in the French region of Occitanie (Barbaro et al., 2022). More precisely, the region sampled is part of the Zone Atelier Pyrénées Garonne (Ouin et al., 2021), considered a long-term socio-ecological research site by French research institute CNRS since 2017. Out of the 79 sampling sites from which recordings were obtained, 60 are found in the Aurignac canton, 16 are found either in alpine meadows or at the edge between alpine meadows and subalpine forests at altitudes between 1400 and 2100 m in the Ariège department, and the remaining 3 are found in lowland agricultural areas in South-West Occitanie (Fig. 1a).

**Table 1:**
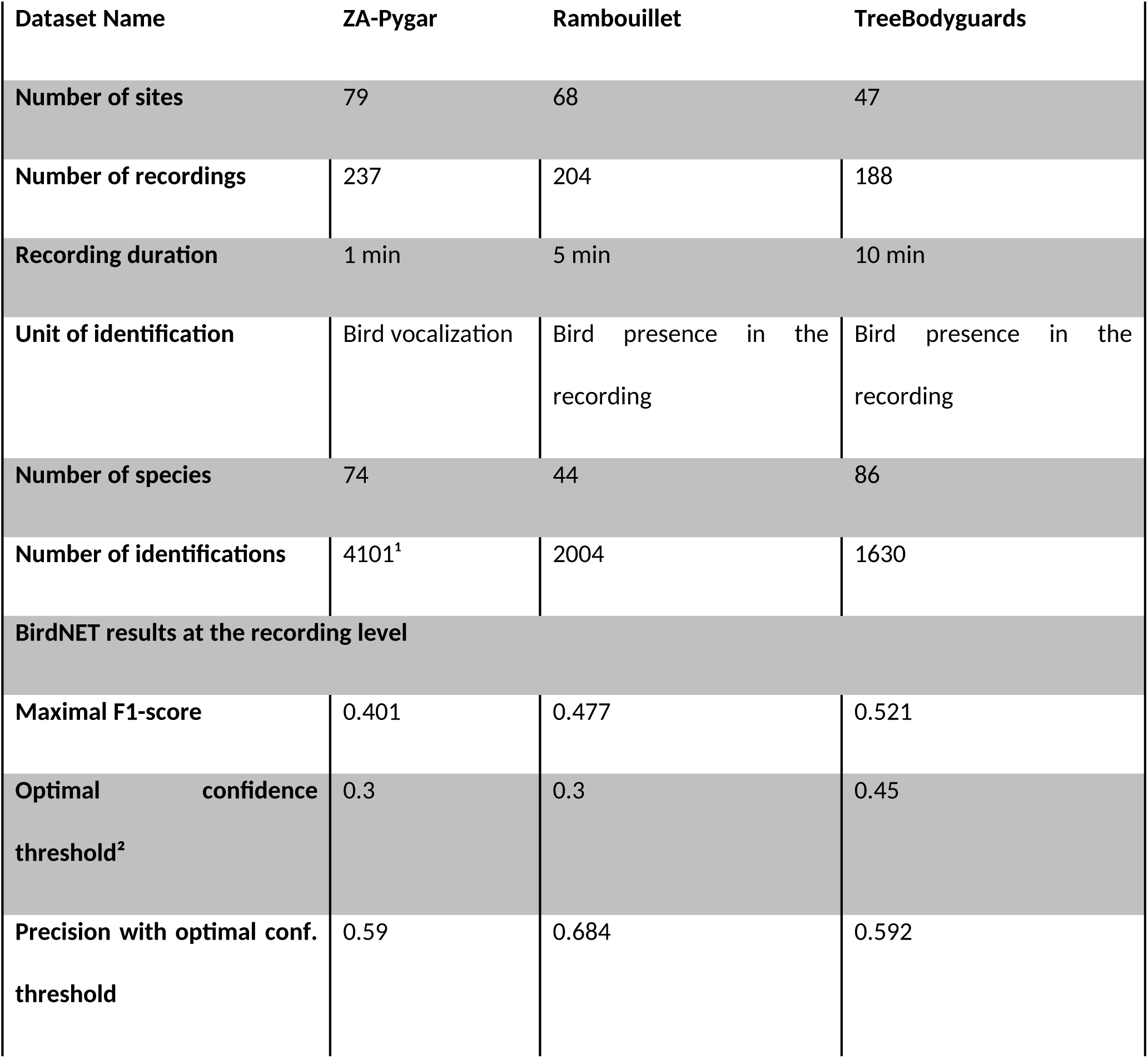
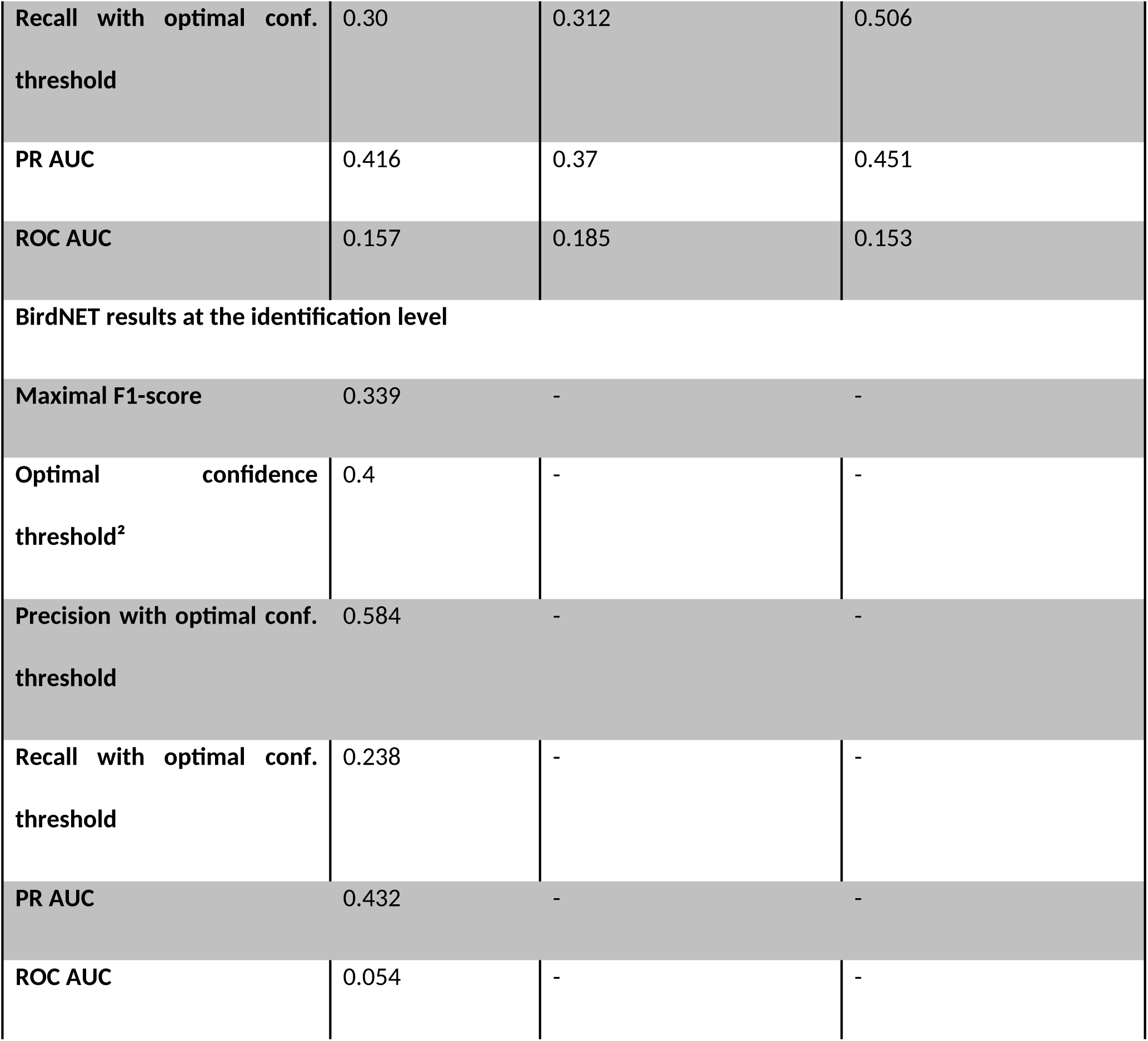
Global description of the three datasets studied along with multiple indicators of BirdNET performance for each dataset. ^1^3340 songs, 754 calls and 7 drummings ^2^Confidence threshold with the highest F1-score

Our second dataset, hereafter referred to as Rambouillet, consists of a total of 204 5-minute recordings collected during the COVID-19 lockdown of 2020 in 68 different sampling sites in the public forest of Rambouillet (Fig. 1b). This lowland sub-Atlantic broadleaf forest, covering 220 km^2^, is located in South-West Île-de-France (France), about 70 km from Paris (Barbaro et al., 2023). Finally, our third dataset, hereafter referred to as TreeBodyguards, was inherited from a pan-European citizen science project and consists of a total of 188 10-minute recordings collected in 47 different sampling sites from 17 European countries ranging from Spain to North-West Russia (Fig. 1c) (Valdés-Correcher et al., 2021).

### 2.2. Manual bird identification by experts

Bird vocalizations have been identified by expert birders in the three datasets. In the ZA-Pygar dataset, all bird sounds found within the first minute of each of the 237 recordings were manually identified and annotated by a single trained birder (DF) by drawing time-frequency bounding boxes around every single vocalization on recording spectrograms (Appendix S2, Fig. S1). In the two other datasets, a list of species present in each recording was generated after the listening, the specific vocalization times of each species remaining unknown. In the Rambouillet dataset, each 5-minute recording selected was manually listened to once, all species detected being listed regardless of their total vocalization time by a single trained birder (LB). As for the TreeBodyguards dataset, given its wide geographical range, we distributed the recordings among 21 expert birders (see Acknowledgements), each of them manually listening to and identifying bird sounds from between 4 (one site) to 52 recordings (13 sites).

### 2.3. BirdNET-assisted analysis of recordings

The BirdNET version chosen for this study is BirdNET-Analyzer v2.4, the most recent release of the algorithm at the time of publication. BirdNET takes as input not only a sound file but also its approximate recording date (in the form of week of the year) and geographic coordinates (Appendix S3). It then splits the input recording into 3-second segments and provides a list of species detected in each segment along with a confidence score assigned to each identification, with values ranging from 0 to 1. By default there is no overlap between consecutive prediction segments, i.e., each segment begins where the previous segment ends, but BirdNET offers the possibility of allowing a certain degree of overlap (Fig. S2). In this study we initially performed the same analyses with allowed overlaps of 0 s, 1 s and 2 s between consecutive segments, and the solidly higher performances obtained with the latter option on the ZA-Pygar dataset (Table 2) made us choose it as the input value to evaluate BirdNET with in all datasets. A detection sensitivity input value of 1.5 was chosen for the analysis of the three datasets for the same reason.

**Table 2:** Area Under the Curve (AUC) scores for both Receiver Operating Characteristic (ROC) and Precision-Recall (PR) curves depending on the detection sensitivity used and the levels of overlap allowed between consecutive prediction segments by BirdNET. The *ID_*, *rec_* and *ds_* prefixes correspond to precision, recall and FPR being calculated at the vocalization/identification level, recording level and dataset level, respectively. Best results are shown in bold and worst in italics. Only results from the ZA-Pygar dataset are included.

| Overlap | Detection sensitivity | ID_PR AUC | ID_ROC<br>AUC | rec_PR<br>AUC | rec_ROC<br>AUC | ds_PR<br>AUC | ds_ROC<br>AUC |
| --- | --- | --- | --- | --- | --- | --- | --- |
| 0s | 0.5 | 0.312 | 0.011 | 0.177 | 0.049 | 0.255 | 0.084 |
| 0s | 1.0 | 0.384 | 0.012 | 0.316 | 0.051 | 0.360 | 0.076 |
| 0s | 1.5 | 0.382 | 0.023 | 0.420 | 0.136 | **0.449** | 0.130 |
| 1s | 0.5 | 0.374 | 0.014 | 0.213 | 0.051 | 0.363 | 0.038 |
| 1s | 1.0 | 0.408 | 0.018 | 0.336 | 0.060 | 0.409 | 0.062 |
| 1s | 1.5 | 0.388 | 0.031 | **0.423** | 0.143 | 0.428 | 0.148 |
| 2s | 0.5 | 0.363 | 0.036 | 0.246 | 0.058 | 0.354 | 0.057 |
| 2s | 1.0 | 0.428 | 0.036 | 0.380 | 0.062 | 0.445 | 0.058 |
| 2s | 1.5 | **0.432** | **0.054** | 0.416 | **0.157** | 0.446 | **0.152** |

### 2.4. Comparing BirdNET and expert identifications

We compared the results obtained with BirdNET against the identifications made by the expert birders in order to evaluate the performance of the algorithm. BirdNET results corresponding to the ZA-Pygar dataset could be analyzed at a higher level of detail than those obtained for the other two datasets, since its temporally annotated identifications make it possible to know the specific vocalization times of each species. For our evaluation we defined 4 possible categories –True Positives (TP), False Positives (FP), True Negatives (TN) and False Negatives (FN)– for each species and acoustic sample, the categorization criteria being slightly variable among the three acoustic datasets analyzed (Table S1).

Based on the categorization described, we calculated the precision, the recall and the False Positive Rate (FPR) of the algorithm for each minimum confidence threshold studied, going from 0.10 to 0.99 with a step of 0.01 (e.g., a minimum confidence threshold of 0.5 implies that all BirdNET identifications with confidence levels lower than 0.5 were filtered out). Precision is a measure of the reliability of BirdNET identifications, i.e., it estimates the probability that a given BirdNET identification will be correct, and is calculated by dividing the number of species correctly detected by the total number of species detected by the algorithm in the acoustic sample analyzed. Recall estimates the probability that a species present in the acoustic sample will be correctly detected by BirdNET, and is calculated by dividing the number of species correctly detected by the total number of species actually present in the acoustic sample analyzed. Finally, FPR estimates the probability that a species absent from the acoustic sample will be detected by BirdNET, and is calculated by dividing the number of species mistakenly detected by the total number of species absent from the acoustic sample analyzed (only considering those featuring in the species lists generated following the procedure explained in Appendix S3). The specific formulas used are the following:

- Precision = TP / (TP + FP)
- Recall = TP / (TP + FN)
- FPR = FP / (FP + TN)

Different conclusions can be drawn from these metrics at different levels of analysis. In the ZA-Pygar dataset, measuring precision at the identification level and recall at the vocalization level can allow us to estimate what percentage of BirdNET identifications will be correct and what percentage of bird vocalizations will be correctly detected by BirdNET, thus providing us with a fine-grained picture of BirdNET performance. However, since the total vocalization time of any given species varies significantly across different recordings, evaluating BirdNET based on the correctness of each individual identification could result in a small number of recordings with long vocalization times having a disproportionate weight over a greater number of recordings with shorter vocalization times. Thus, calculating precision and recall at the recording level can be an appropriate way of homogenizing the weight of each recording in the final results. Finally, calculating precision and recall at the level of the whole acoustic dataset can inform us about the reliability and exhaustiveness of BirdNET results when using them to provide us with a broad picture of the composition of a bird community recorded over multiple hours or days.

We therefore calculated precision, recall and FPR at all three different levels of analysis: (i) at the identification/vocalization level (only for the ZA-Pygar dataset), (ii) at the recording level and (iii) at the dataset level. The metrics calculated at the first level will be referred to as ID_precision, ID_recall and ID_FPR, the metrics calculated at the second level will be referred to as rec_precision, rec_recall and rec_FPR and the metrics calculated at the third level will be referred to as ds_precision, ds_recall and ds_FPR (Table 3). It is important to note that, since our categorization criteria imply that one correct identification can override any number of incorrect identifications of the same species within the same acoustic sample, the values obtained for these metrics will be highly influenced by the level of analysis chosen. Longer acoustic samples provide more opportunities for BirdNET to detect a given species, either correctly or incorrectly, but the asymmetric weight given to correct identifications over incorrect ones implies that the number of TPs will scale more rapidly with time than the number of FPs. This results in a bias towards more positive results when longer acoustic samples are used.

**Table 3:** Description of the different metrics used to evaluate BirdNET performance. ^1^Only calculated for the ZA-Pygar dataset ^2^Only considering those featuring in the species lists generated following the procedure described in Appendix S3 ^3^Corresponding to 1 minute in the ZA-Pygar dataset, 5 minutes in the Rambouillet dataset and 10 minutes in the TreeBodyguards dataset

| **Metric** | **Acoustic sample** | **Description** |
| --- | --- | --- |
| **ID_precision<sup>1</sup>** | BirdNET identification | Proportion of BirdNET identifications that are correct (match in time and species with an annotation by an expert). |
| **ID_recall<sup>1</sup>** | Manually annotated bird vocalization | Proportion of manually annotated bird vocalizations that have been correctly identified by BirdNET |
| **ID_FPR<sup>1</sup>** | BirdNET prediction segment | Number of species mistakenly detected by BirdNET as a fraction of all species actually absent from the prediction segment<sup>2</sup> |
| **rec_precision** | Entire recording<sup>3</sup> | Proportion of species detected by BirdNET that are actually present in the recording |
| **rec_recall** | Entire recording<sup>3</sup> | Proportion of species present in the recording that have been correctly detected by BirdNET |
| **rec_FPR** | Entire recording<sup>3</sup> | Number of species mistakenly detected by BirdNET as a fraction of all species actually absent from the recording<sup>2</sup> |
| **ds_precision** | Acoustic dataset | Proportion of species detected by BirdNET that are actually present in the acoustic dataset |
| **ds_recall** | Acoustic dataset | Proportion of species present in the acoustic dataset that have been correctly detected by BirdNET |
| ds_FPR | Acoustic dataset | Number of species mistakenly detected by BirdNET as a fraction of all species actually absent from the acoustic dataset<sup>2</sup> |

Once calculated, we used these values to plot the Receiver Operating Characteristic (ROC) and Precision-Recall (PR) curves, both with an estimation of the Area Under the Curve (AUC) (Davis & Goadrich, 2006). The ROC curve consists in plotting recall against FPR for each minimum confidence threshold studied in order to show the trade-off between the two metrics: as the minimum confidence score required to accept BirdNET identifications increases, the number of FPs will decrease, but so will the number of TPs. The PR curve, on the other hand, plots precision against recall for each minimum confidence threshold used, showing the trade-off between these two metrics as well. The AUC is, in both cases, a measure of the predictive power of the algorithm with a range of possible values going from 0 to 1, higher values being indicative of a higher predictive power.

Another metric that we used to evaluate the performance of the algorithm for each minimum confidence threshold studied is the F-score. The F-score measures the overall predictive power of the algorithm by considering both precision and recall scores, the weight assigned to each variable depending on the β coefficient:

F-score = (β^2^ + 1) * precision * recall / (β^2^ * precision + recall)

An F-score with β = 1 assigns the same importance to precision and recall, whereas an F-score with β > 1 weighs recall more heavily than precision, and vice versa for β < 1. We performed F-score analyses using three different β values: β = 1 as a standard value to facilitate the comparison of our results with those of other studies, β = 0.25 to reflect a clear prioritization of precision over recall and β = 0.1 to reflect an even more marked asymmetry between the importance assigned to these two metrics. We chose such asymmetrical coefficients because we estimate that ensuring high precision levels is paramount in most biodiversity research projects, i.e., it is usually considered preferable not to detect present species rather than mistakenly detecting non-present ones (Tolkova et al., 2021).

### 2.5. Factors influencing BirdNET performance

Using only the Za-Pygar dataset, we further analyzed how BirdNET performance might be influenced by factors related to the acoustic dataset analyzed (total duration recorded, habitat recorded, passive recorder used, preponderance of biophony over anthropophony and geophony (Bradfer-Lawrence et al., 2023; see Appendix S6), and number of species vocalizing at the same time) as well as the number of recordings available online for each species as a proxy for the size of the training dataset used by BirdNET.

To estimate the influence of the size of the dataset (i.e., total recording duration) analyzed on birdNET performance, we calculated recall and FPR at the dataset level (ds_recall and ds_FPR) for 23 different subset sizes going from 10 to 230 recordings using a step increase of 10. For each subset size, we performed 200 random selections of recordings from the whole set of 237 recordings making up the ZA-Pygar dataset and then averaged out their results (i.e., we generated 200 random subsets of 10 recordings and then averaged out the ds_recall and ds_FPR corresponding to each of these subsets; likewise for all other subset sizes until 230). This number of random selections proved to be sufficient for results to be solidly consistent across multiple iterations of the same analysis.

As for the variability of the predictive power of BirdNET across species, we distinguished between two different types of causal factors: (i) factors related to the difficulty inherent in identifying certain vocalizations, such as some acoustic patterns being easier to identify than others, or bird species with vocalizations highly similar to those of other species being harder to identify than species with more idiosyncratic sounds, and (ii) the size and quality of the acoustic data available for each species on the online platforms used as sources of data for the training of the algorithm. The first type of factor being considered out of scope for this study, we analyzed the influence of species-specific data availability on BirdNET performance. Even though we do not have direct access to the training datasets used by BirdNET, we know that the Xeno-canto platform (Xeno-canto, 2023) and the Macaulay Library of Natural Sounds (Macaulay, 2023) were used as sources of audio data for this purpose (Kahl et al., 2021). Thus, we used the total number of foreground recordings available on both platforms for each species analyzed as a rough proxy for the species-specific availability of recordings during the training phase of the algorithm. More specifically, we examined the correlation between the number of recordings available on these platforms and BirdNET precision, recall and F1-scores across species. This allowed us to roughly estimate the explanatory power of species-specific recording availability over the variability of BirdNET performance across species. Species detected fewer than 10 times were filtered out from the analysis, their sample sizes being too low for results to be minimally reliable. Finally, it is important to note that, unless explicitly specified, all Fig.Figs and statistical results correspond to the analysis of recordings with BirdNET-Analyzer v2.4, using a detection sensitivity of 1.5 and an overlap window of 2 s between consecutive prediction segments.

## 3. RESULTS

### 3.1. Optimizing BirdNET parameters

Our evaluation of BirdNET performance under different input parameter configurations suggests that 1s overlap windows improve AUC scores on both ROC and PR curves with respect to the default 0-second overlap, and that 2s overlaps improve these scores even further (Table 2). This improvement seems to arise from the fact that high degrees of overlap facilitate the capture of a substantial part of a given bird vocalization within a single prediction segment (Fig. S1). Since longer vocalization times within a given prediction segment are predictive of higher ID_recall levels (Fig. S3; Spearman, r_s_ = 0.956 and p < .001), an increase in overlap between consecutive prediction segments should correspondingly result in higher recall scores. Likewise, the predictive power of BirdNET as measured by the ROC and PR AUC scores seems to improve along with higher values of the detection sensitivity parameter (Table 2), suggesting that a detection sensitivity of 1.5 (from a range of possible values going from 0.5 to 1.5) might be the optimal value for overall performance.

### 3.2. Assessing BirdNET performance

We found the confidence scores provided by the algorithm to be positively correlated with the actual probability that the corresponding identification is correct (Figs 2 and S4; Kruskal-Wallis, H(1) = 5977.7 and p < .001). This means that a minimum confidence threshold can be established so that identifications with low confidence scores are filtered out, thus retaining only the most reliable ones. In accordance with this premise, the ROC and PR curves in Fig. 3 reveal a clear trade-off between the precision and recall of the algorithm, as well as between recall and FPR: the higher the minimum confidence score required to accept BirdNET identifications, the higher the precision and the lower the FPR will be. Nonetheless, improvements in these two metrics have to be weighed against the lower recall associated with a higher level of selectiveness. Average precision scores by species improve considerably by setting higher minimum confidence thresholds (Fig. S5), recall scores by species following the opposite trend (Fig. S6).

**Fig. 2:**
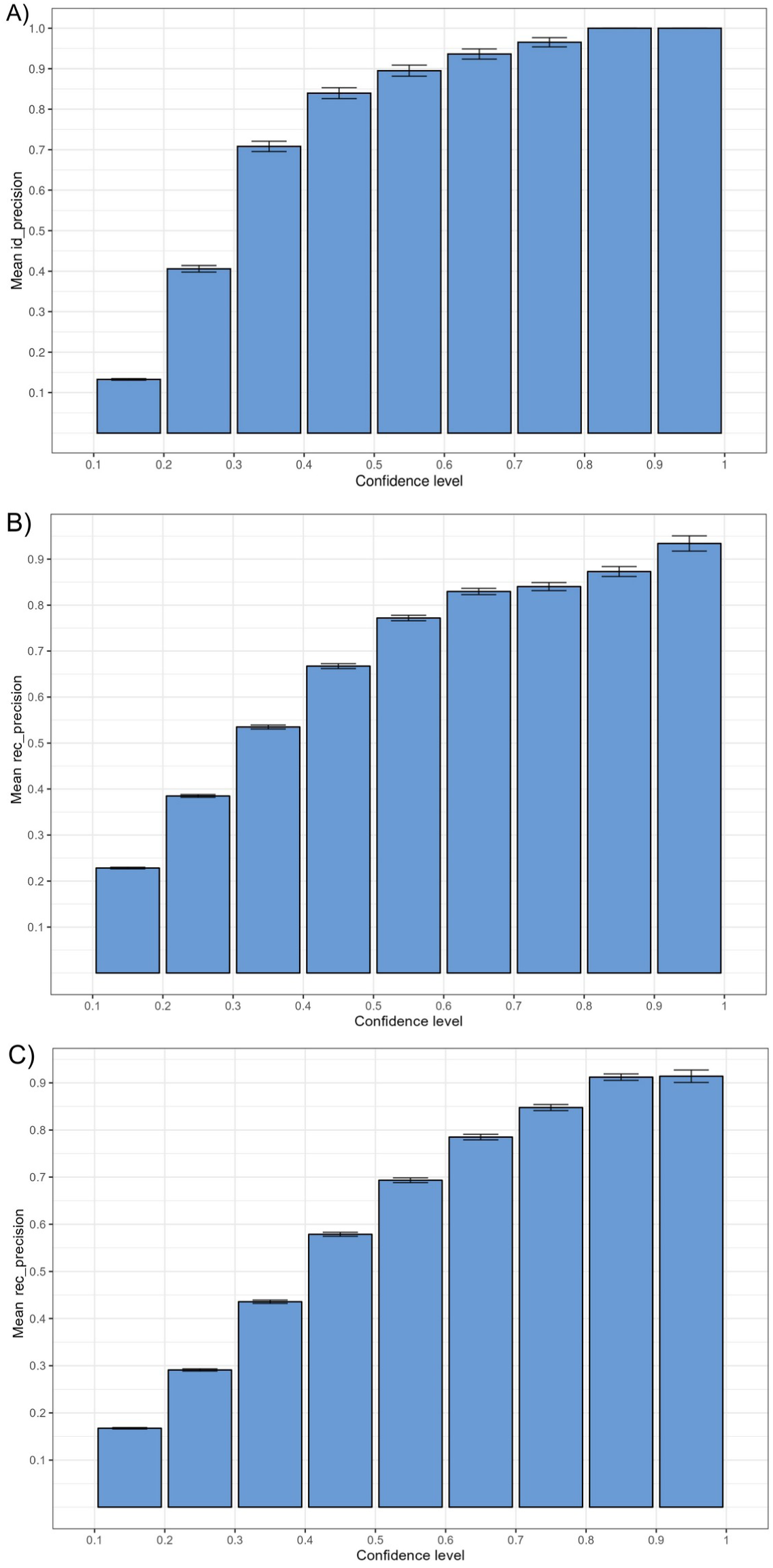
Mean precision, along with the standard error bar, of BirdNET identifications for different ranges of confidence scores. Results correspond to (A) ID_precision in the ZA-Pygar dataset, (B) rec_precision in the Rambouillet dataset, and (C) rec_precision in the TreeBodyguards dataset.

**Fig. 3:**
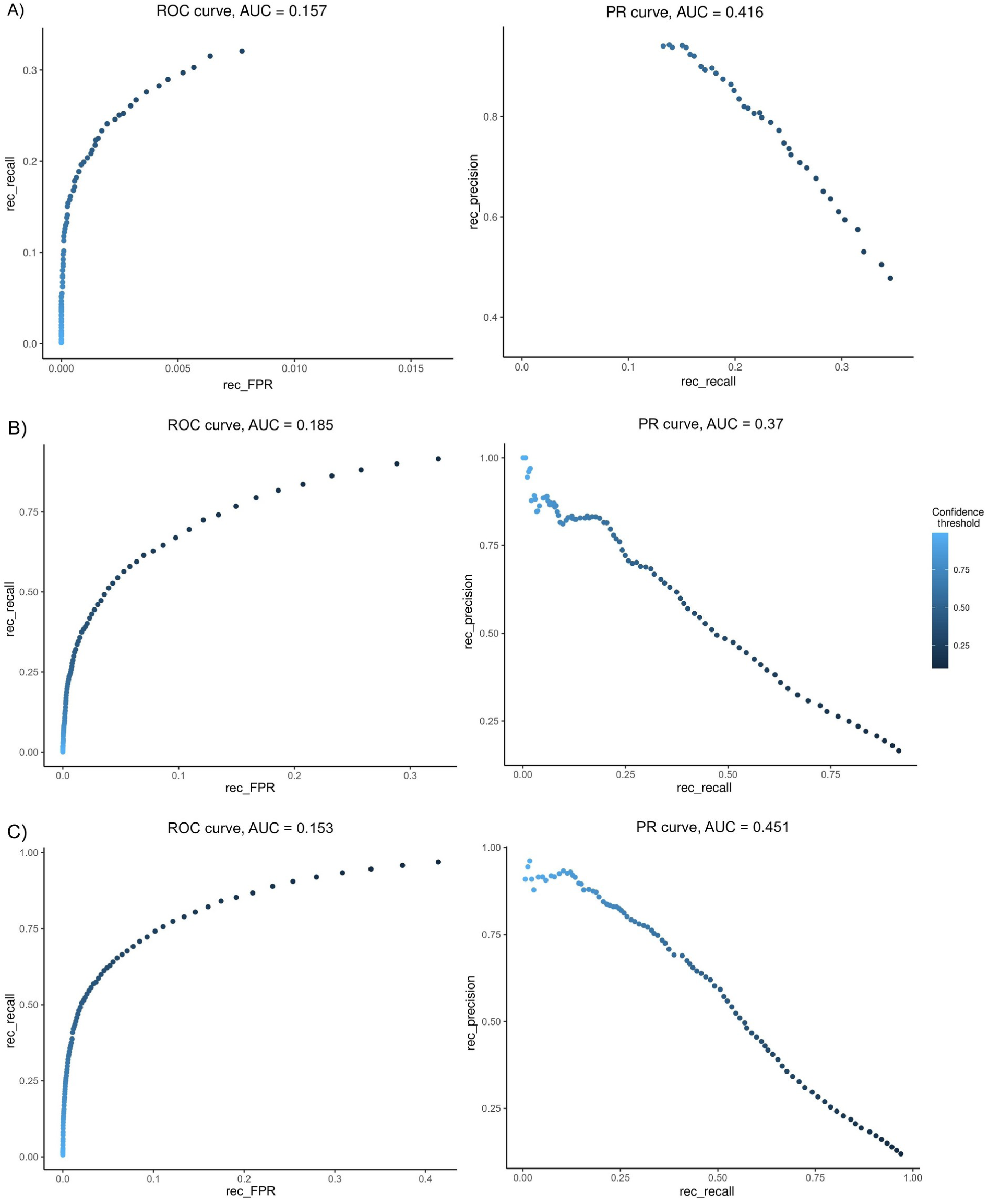
Receiver Operating Characteristic (left) and Precision-Recall (right) curves of BirdNET-Analyzer. The three plots correspond to results obtained at the recording level for the (A) ZA-Pygar, (B) Rambouillet and (C) TreeBodyguards datasets.

When analyzing BirdNET results at the recording level for the ZA-Pygar dataset, we found that the optimal minimum confidence thresholds for maximizing global F-scores for β-values of 1, 0.25 and 0.1 are 0.3, 0.55 and 0.65, respectively (Fig. 4). The results obtained for each of these thresholds show that, as the minimum confidence score required to accept BirdNET identifications becomes stricter, both TPs and FPs decrease while FNs rise sharply (Fig. S7). This causes both the number of species correctly detected and the number of species mistakenly detected to vary considerably depending on the minimum confidence threshold used. While a minimum confidence threshold of 0.3 allows for a ds_recall of 0.65 (i.e., the correct detection of 65% of the species present in the dataset), a threshold of 0.55 reduces this Fig. to 0.55 and a threshold of 0.65 reduces it further to 0.5. On the other hand, ds_FPR (i.e., the proportion of species absent from the dataset that are mistakenly detected by the algorithm) varies even more strongly along with the confidence threshold used, with respective values of 0.41, 0.07 and 0.03. In absolute terms, this translates into 48, 41 and 37 species correctly dectected and 85, 15 and 6 species mistakenly detected, respectively.

**Fig. 4:**
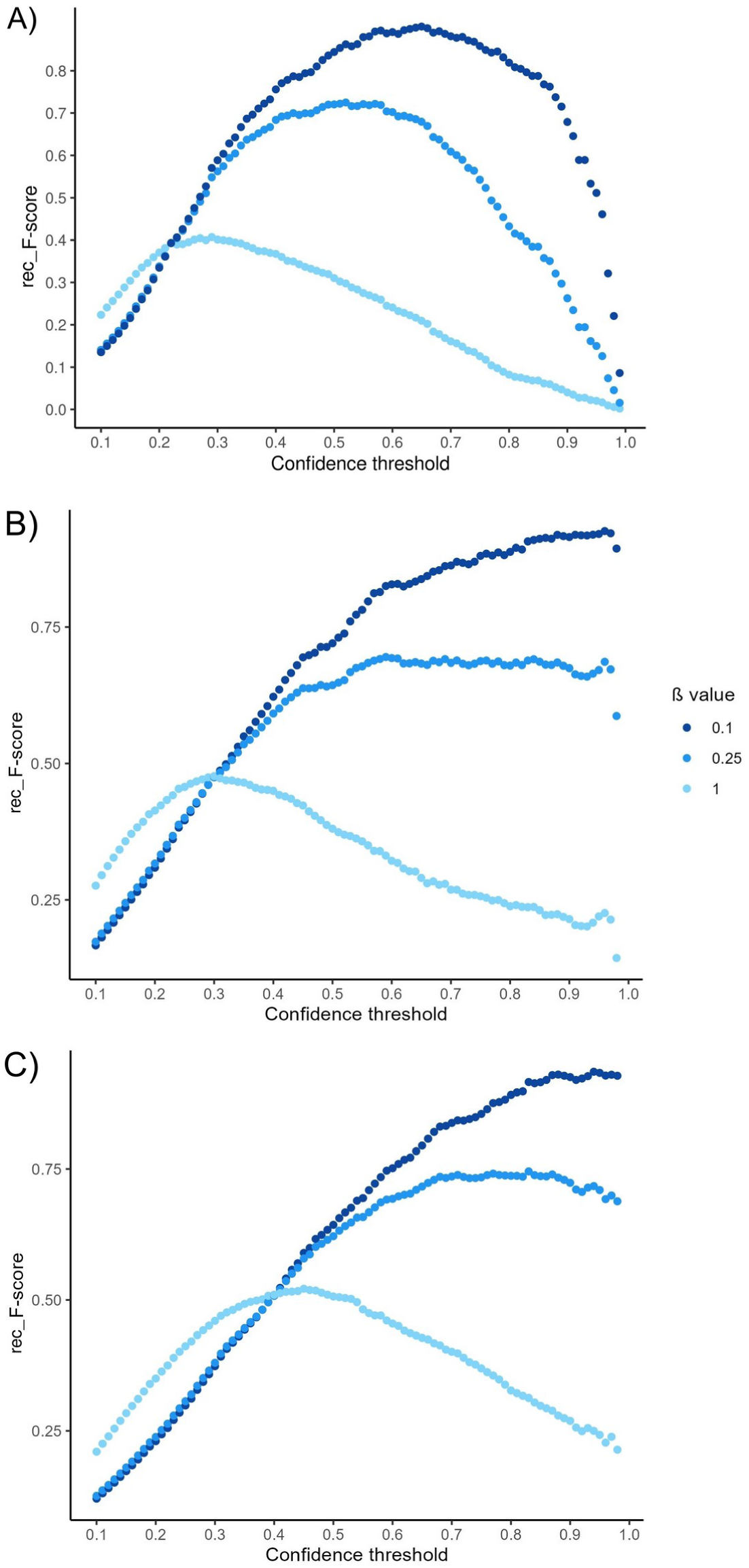
F-score curves of BirdNET-Analyzer for β values of 0.1, 0.25 and 1. The three plots correspond to results obtained at the recording level for the (A) ZA-Pygar, (B) Rambouillet and (C) TreeBodyguards datasets.

It is important to note that a non-negligible number of species present in the recordings have only been manually detected a small number of times (Fig. S8). Hence, we deem the estimated predictive power of the algorithm for these species to be highly unreliable. The difference between Figs S7a and S7d suggests that limiting the analysis to species detected –either by BirdNET or by the expert birder– in a minimum of 10 recordings, while filtering out less common species, results in a substantially higher overall performance of the algorithm. More specifically, when analyzing the ZA-Pygar dataset with a confidence threshold of 0.3, this filtering procedure improves rec_precision from 0.59 to 0.71 without compromising rec_recall, which actually remains stable at 0.32. The results obtained after applying this filter (Fig. S7d) also suggest that the variability in BirdNET predictive power across species is not as pronounced as what we might infer from Fig. S7a, since the low number of data points available for many species leads to more variable outcomes.

Overall, our analyses yielded highly consistent results across the three datasets studied, with maximal F1-scores (corresponding to minimum confidence thresholds between 0.3 and 0.45) ranging from 0.4 to 0.52 at the recording level (Table 1). This roughly means that, in the best case, we can expect two errors (FP or FN) for each correct BirdNET prediction. BirdNET performance is lower (maximal F1-score of 0.33 for a minimum confidence threshold of 0.4) when results are analyzed at the identification/vocalization level (i.e., 1 correct prediction for every 3 errors).

### 3.3. Factors influencing BirdNET performance

In order to properly isolate the variables of interest and ensure that the heterogenous identification procedures and recording durations used in the three datasets do not have a confounding effect, only recordings from the ZA-Pygar dataset are included in the following analyses.

Regarding the influence of sample size on BirdNET performance, there appears to be a positive linear correlation between ds_F1-scores and the logarithm of the number of recordings sampled (Fig. 5; linear regression, adjusted R^2^ = .539, p < .001). The same seems to be true for ds_TPR (adjusted R^2^ = .376, p < .001) and ds_recall (adjusted R^2^ = .095, p < .001) as well, in all cases after controlling for the minimum confidence threshold used. As can be observed in Fig. 5, ds_FPR shows a pronounced decrease when higher confidence scores are required. ds_recall, while following the same trend, shows differences of more moderate proportions as a response to increased minimum confidence thresholds. This suggests that a given global recall target can be reached either by setting a low enough minimum confidence threshold or by setting a high minimum confidence threshold and procuring a large enough acoustic dataset, the latter approach resulting in substantially lower ds_FPR values.

**Fig. 5:**
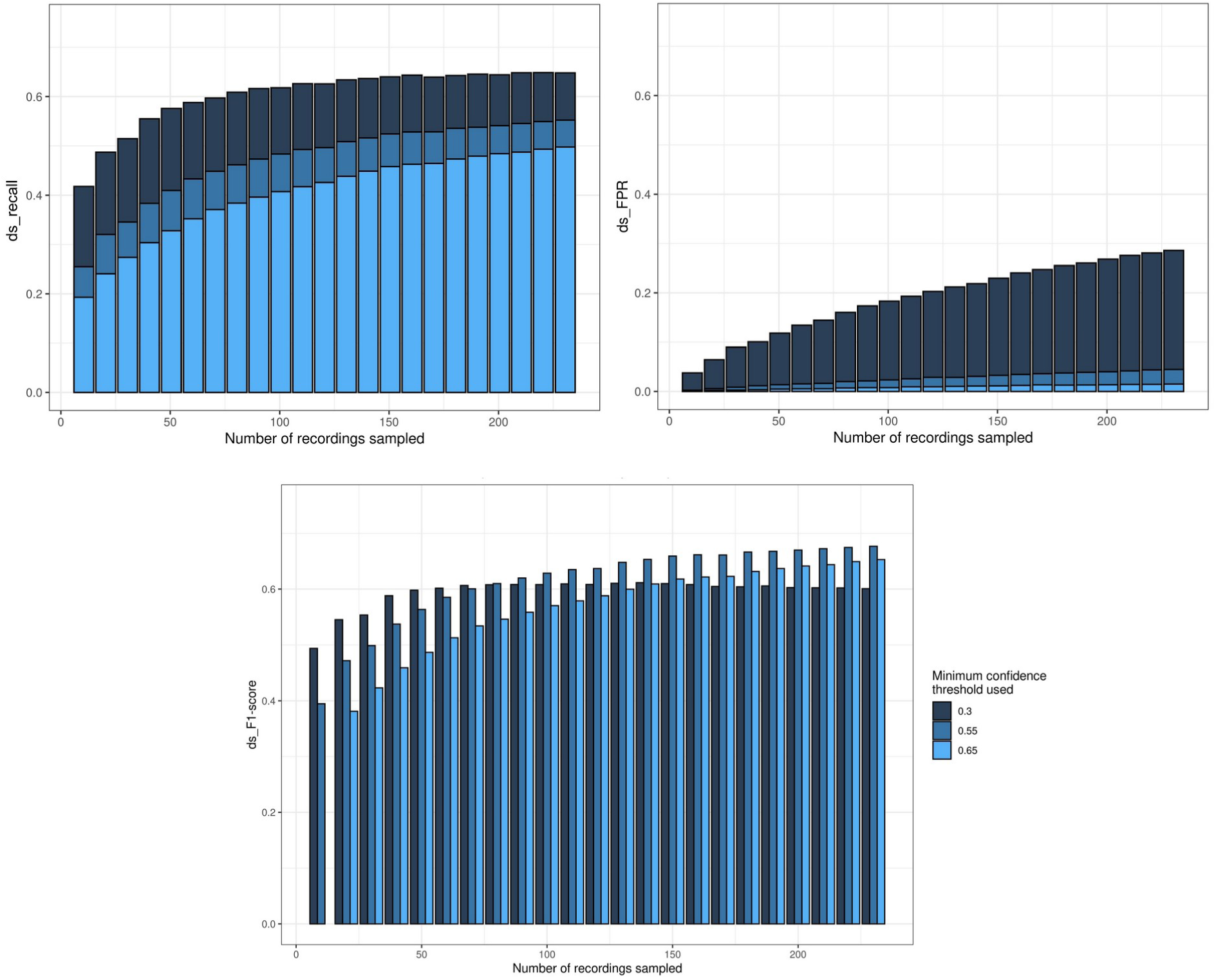
Dataset performance depending on the number of recordings included in the analysis. ds_recall (top-left), ds_FPR (top-right) and ds_F1 scores (bottom) are shown for virtual datasets of increasing size depending on the minimum confidence threshold required to validate BirdNET identifications. Only recordings from the ZA-Pygar dataset are included.

Other analyses performed suggest that the number of different bird species vocalizing simultaneously seems to have a significant influence on ID_precision (Fig. S9; Spearman, r_s_ = 0.122 and p < .001) but not on ID_recall and ID_F1-scores (Spearman, r_s_ = −0.051 and p = .285 for ID_recall, r_s_ = 0.017 and p = 0.72 for ID_F1-score). In contrast, neither the recorder used nor the predominant habitat sampled seem to affect BirdNET performance (Fig. S10), since neither rec_precision, rec_recall nor rec_F1-scores show significant differences across different habitats (Kruskal-Wallis, H_4_ = 1.11 and p = .775 for rec_precision, H_4_ = 0.79 and p = .852 for rec_recall and H_4_ = 1.24 and p = .744 for rec_F1_score) or recorders (Kruskal-Wallis, H_3_ = 4.08 and p = .130 for rec_precision, H_3_ = 8.92 and p = 0.116 for rec_recall and H_3_ = 2.04 and p = 0.36 for rec_F1_score). Finally, the number of foreground recordings available on Xeno-canto and the Macaulay Library for each species studied appears to have a significant influence on species-specific rec_precision, rec_recall and rec_F1-scores. More concretely, the logarithm of the number of recordings available seems to correlate positively with rec_precision and rec_F1-score but negatively with rec_recall (Fig. 6; Spearman, r_s_ = 0.506 and p < .001 for rec_precision, r_s_ = −0.484 and p = .002 for rec_recall and r_s_ = 0.599 and p < .001 for rec_F1-score).

**Fig. 6:**
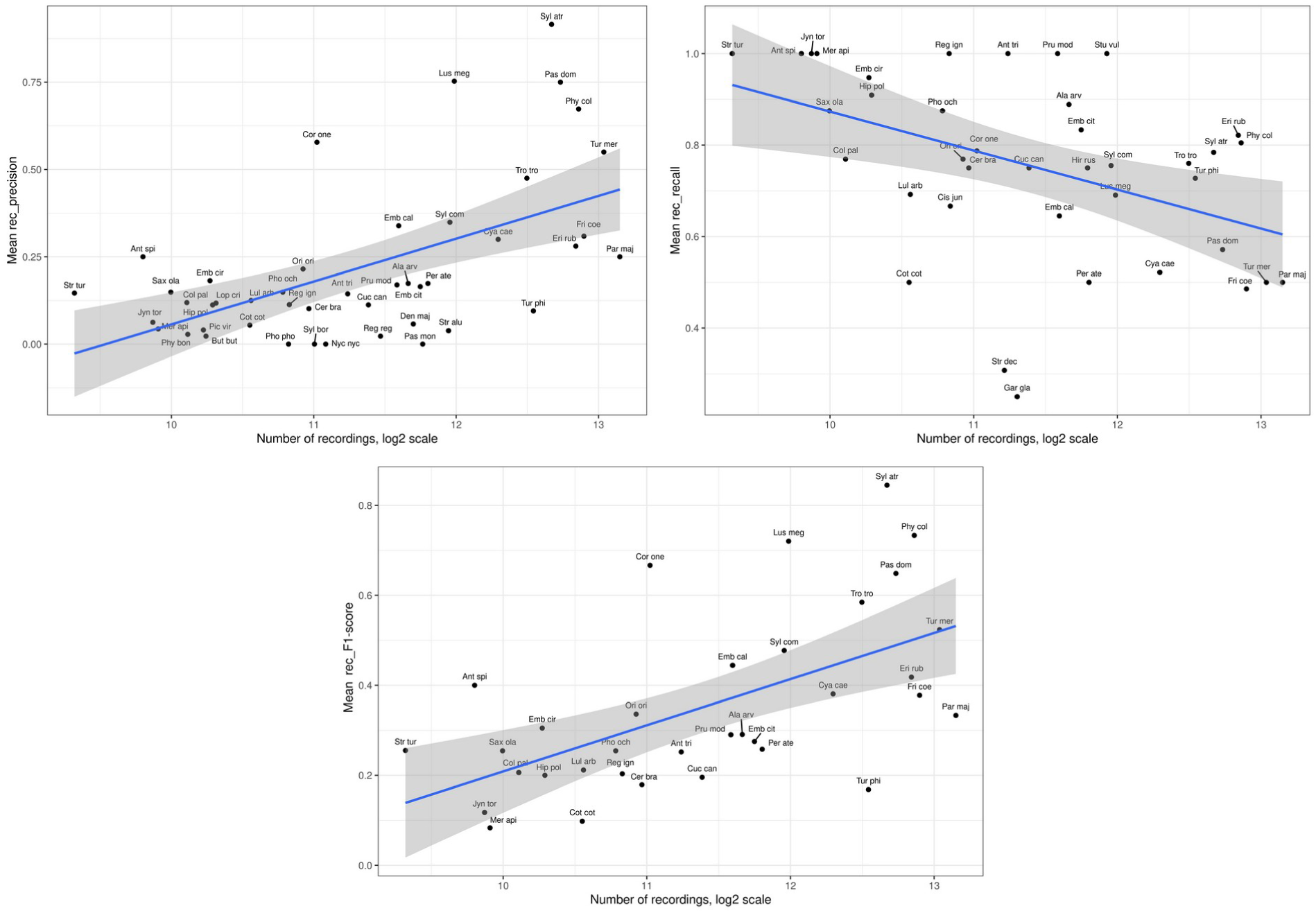
Relationship between BirdNET performance and the number of foreground recordings available for each species on the Xeno-canto and Macaulay Library platforms. More specifically, the mean rec_precision (top-left), rec_recall (top-right) and rec_F1 (bottom) scores obtained for each species are plotted against the base-2 logarithm of the number of recordings available online for the species in question. Only species having been detected 10 or more times by BirdNET are included in the precision analysis, only species with at least 30 s of (manually annotated) total vocalization time are included in the recall analysis and only species meeting both criteria are included in the F1-score analysis. The minimum confidence threshold used is 0.1, and only recordings from the ZA-Pygar dataset are included.

## 4. DISCUSSION

Even with its current limitations, BirdNET appears as a promising tool to assist in the assessment of bird community composition through the automated processing of large-scale ecoacoustic data. Our analyses reveal that BirdNET can provide us with reasonably high levels of precision or recall, at least in the regions studied, but the inevitable trade-off between these two metrics prevents satisfactory results for both at the same time (maximal F1-score <0.5). Admittedly, the current recall of the algorithm when adjusted to ensure high levels of precision is markedly low. Despite BirdNET still having considerable room for improvement in both precision and recall scores, it is important to note that identifications by field observers are not exempt from errors either. Recent studies suggest that acoustic identifications by highly experienced birders present precision scores no higher than 0.94 and recall scores no higher than 0.89 (Farmer et al., 2012; Campbell & Francis, 2011), implying that BirdNET identifications with exceptionally high confidence scores might prove to be as accurate or more as those made by reasonably experienced birders.

Regarding precision scores in particular, our results are in line with previous studies (Sethi et al., 2021; Cole et al., 2022) showing that BirdNET identifications can be highly reliable, especially for common species (Fig. 6), provided that a sufficiently high minimum confidence threshold is used (Fig. S5). Conveniently, confidence scores assigned to each identification by the algorithm show a high degree of correspondence with the actual probability that the identification in question is correct (Fig. 2). This makes it possible to reliably adjust the precision of the identifications to the specific requirements of each research project using BirdNET.

Our results further suggest that BirdNET precision scores can be improved not only by raising the minimum confidence threshold used, but also by filtering out species not having reached a minimum number of BirdNET detections across the whole dataset (Fig. S5). We found, however, that species such as garden warblers (*Sylvia borin*) and Western Bonelli’s Warblers (*Phylloscopus bonelli*) were identified by BirdNET on a high number of occasions, always incorrectly, and exceeded our minimum occurrence frequency threshold (Fig. S7d). Non-negligible numbers of FPs for other non-present species suggest caution around their identifications as well. On the positive side, these misidentifications seem to be effectively filtered out by establishing strict enough minimum confidence thresholds (Fig. S7c). It is important to note, though, that background sounds can vary considerably between different locations and times of the year, so the list of problematic species can turn out to be substantially different in other spatio-temporal contexts. We therefore recommend that, prior to its use, a preliminary assessment of BirdNET be conducted with soundscapes from the target study site. This would allow researchers to preemptively identify the species appearing most often as FPs and to pay special attention to identifications of these species when examining BirdNET results. Another possible method to reduce FPs, as suggested by a recent study (Toenies & Rich, 2021), would be to filter out BirdNET identifications of diurnal species when analyzing nocturnal recordings, thus effectively preventing most dog barks and frog croaks from being misclassified as diurnal waterbird calls.

Our results also make it clear that BirdNET performance shows a high degree of variability across different species (Figs S5, S6, S7). However, these results do not provide information on the degree to which this variability is due to (i) cross-species differences in the amount or quality of recordings available to train the algorithm with, (ii) the inherent difficulty in identifying vocalizations with certain acoustic patterns, or (iii) the identification difficulty arising from two or more species emitting highly similar sounds. Assuming that the first obstacle is the most tractable among the three mentioned, the higher the percentage of cross-species variation in BirdNET performance that is explained by recording availability, the more optimistic we should be regarding the improvement of the performance of the algorithm over time. In this regard, our results suggesting that 19% of the variance in F1-scores between species can be explained by online recording availability (Fig. 6) provide ground for optimism about the future reliability of BirdNET. More concretely, this could mean that the diagnostic capacity of BirdNET for less common species could be considerably improved by enlarging the pool of available acoustic data for these species.

We presume that a higher availability of acoustic data could facilitate BirdNET performance through two different mechanisms: (i) by enabling BirdNET to enlarge the size of its training dataset, and (ii) by improving the quality of the recordings comprising its training dataset. Acoustic quality scores are included in the metadata of recordings on Xeno-canto and the Macaulay Library, and higher-quality recordings were prioritized in the selection of training data for the BirdNET algorithm (Kahl et al., 2021). Hence, we would expect the average acoustic quality to be higher in the training datasets of species with larger pools of available recordings to choose from. Insomuch as both higher-quality recordings and larger training datasets could facilitate a better adjustment of the algorithm to the target bird vocalizations, these factors could provide a plausible explanation to the positive correlation observed between species-specific acoustic data availability and BirdNET predictive power.

We further found a significant linear correlation between the logarithm of the recording time analyzed and the proportion of recorded species having been correctly detected by BirdNET (Fig. 5). A possible explanation to this correlation would be that the short total vocalization time recorded for the least frequent species often results in no correct detections of these species by BirdNET. Thus, insofar as longer recording times contain a greater number of vocalizations per species in expectation, we should expect larger acoustic datasets to provide more chances for the algorithm to correctly detect less frequent species. We therefore hypothesize that a sufficiently high number of recording hours analyzed could partially compensate for the low recall scores obtained when high minimum confidence thresholds are used. Hence, were the necessary conditions to be met, BirdNET might prove successful in capturing most bird species present in a study site while also ensuring a relatively low number of FPs. These results, in line with recent findings (Toenies & Rich, 2021; Cole et al., 2022; Wood et al., 2021), are encouraging with respect to the potential of BirdNET to reliably and comprehensively describe bird communities in research projects with large amounts of ecoacoustic data at their disposal.

Our results also show that the predominance of biophony over anthropophony and geophony in recordings, as measured by NDSI (Kasten et al., 2012), appears to be positively correlated with recall scores (see Appendix S6, Fig. S11). We propose three possible hypotheses explaining this correlation: (i) anthropogenic and geological sounds might hinder the performance of the algorithm by concealing or blurring bird vocalizations, (ii) BirdNET might have mostly been trained with recordings with relatively low levels of background noise, thus being less well calibrated to identify bird vocalizations in noisy recordings, and (iii) high NDSI scores can be indicative of high-volume bird vocalizations, which might to be easier to identify than low-volume bird vocalizations (Pérez Granados, 2023). If the second hypothesis were correct, we consider it plausible that the application of a noise reduction filter to recordings prior to their analysis with BirdNET could result in an enhanced performance of the algorithm.

Overall, the results of this study may provide valuable information for research groups considering the possibility of using BirdNET for the automated analysis of passively-collected environmental soundscapes. However, these results provide an overview of the current potential of BirdNET in a very specific context, and therefore cannot be reliably extrapolated to other regions with different habitats, levels of anthropogenic pressure or bird community compositions. Care must also be taken when extrapolating our results to other areas with similar but geographically distant bird communities, as bird songs and calls can present subtle variations across different regions (Slabbekoorn & Smith, 2002).

Another point that should be borne in mind is that this study starts from the fundamental assumption that all our expert identifications are correct, which is why we used them as the reference to compare BirdNET against. As already mentioned, even highly experienced birders seem to fall short of infallibility (Farmer et al., 2012; Campbell & Francis, 2011), so the possibility that some of the expert identifications included in this study are incorrect cannot be entirely ruled out. As a concluding note, in light of the improvement detected between the recall scores of BirdNET-Analyzer v2.4 and those of older versions of the algorithm –BirdNET-Lite and BirdNET-Analyzer v2.1 to 2.3– that we previously tested with the same data, we consider it plausible that future releases will be capable of identifying bird vocalizations even more reliably and exhaustively. Thus, our results should not be assumed to accurately estimate the performance of future versions of BirdNET, but they should rather be interpreted as a lower bound for their potential.

## Author contributions

Based on an initial idea by MC, he and DF conceived the study, with inputs from LS, BC and LB. LS and BC gathered the TreeBodyguards dataset and LB recorded the ZA-Pygar and Rambouillet datasets. DF performed manual and automated bird identifications and manual annotated recordings for the ZA-Pygar dataset. LB performed manual bird identifications for the Rambouillet dataset. DF and LS performed the computations and data analyses, with inputs from MC and BC. DF wrote the first draft of the manuscript and all authors contributed to shape the final version.

## Acknowledgements

We thank Arndt Hampe, Maarten de Groot, Thomas Boivin, Vojtech Lanta, Lyudmila Pukinskaya, Frédéric Archaux, Thomas Perot, Márton Molnár, Nickolay Sedikhin, Valentin Moser, Elina Mäntylä, Jan Grünwald, Karthik Thrikkadeeri, Anders Mårell, Mona C Bjørn, Becky Thomas, Andreas Prinzing and Lucian Grosu for manual bird identifications of the TreeBodyguards European dataset. We are grateful to Michel Beal, Jean-Luc Témoin, Valérie Delage and Laurent Tillon from Office National des Forêts for their help while recording the Rambouillet dataset. We also thank Thierry Feuillet, Amandine Gasc, Jérémy Froidevaux, Kevin Darras, Yves Bas, Elena Valdés-Correcher, Thomas Delattre for helpful advice and support. We are also indebted to the CESAB Acoucene working group for fruitful discussions.

## Data availability statement

The code used for the analyses performed in this study, as well as both BirdNET and manual identifications/annotations for the three acoustic datasets studied, can be found in the following GitHub repository: https://github.com/dfpl1/BirdNET_study_Occitanie_2023/. The raw acoustic data analyzed is available upon request.

