## Supplementary Information for "Assessing the potential of BirdNET to infer European bird communities from large-scale ecoacoustic data"

**Appendix S1. Acoustic data collection**

***ZA Pygar***

The most dominant study area in this acoustic dataset is the Aurignac canton, covering roughly 190 km² between the Garonne and Gers rivers in South-West France (N 43°16’28 E 0°51’11, WGS 1984). The region falls within an Atlantic subclimate subject to Mediterranean influences, elevation ranging from 200 to 400 m above sea level. Habitats found in the study area include agricultural crops (primarily cereals, sunflower, rapeseed, alfalfa and field bean), mesohygrophilous meadows, grazed and mown grasslands, dry calcareous grass-scrubland mosaics, woodlands and hedgerows. Woodlands are generally fragmented and dominated by Quercus robur and Q. pubescens, although some large forest patches still remain. Hedgerows are composed of various shrubs and mature trees, mainly Quercus spp. Ecoacoustic monitoring is being conducted in Aurignac as part of a multidisciplinary research project started in 1981 with the aim of studying changes in agricultural systems along with their impact on biodiversity and landscape dynamics (Gaüzère et al., 2020).

The 79 sampled sites included in the ZA-Pygar dataset are distributed as follows (Prosbt et al., 2017): 15 are found in farmlands, 18 in mixed farmland- grassland mosaics, 18 in permanent grasslands, 12 in woodlands and 16 in alpine meadows (Fig. 1a). This distribution allows for a fairly representative picture of the composition of breeding bird communities inhabiting the region, and it provides us with a relatively wide range of bird species to test BirdNET with.

Recording procedures vary across the different study areas included in this dataset. In Aurignac we installed Song Meter 4 (Wildlife Acoustics) and AudioMoth (Open Acoustic Devices) acoustic recorders on trees, leaving a minimal distance of 1 Km between any two recorders. We configured both Song Meter 4 and AudioMoth recorders to record with internal omnidirectional microphones at a sampling rate of 24 KHz, allowing us to capture the vast majority of sounds emitted by birds, anourans, cicadas and diurnal orthopterans. Soundscapes were recorded following a discontinuous schedule consisting in recording 30 minutes per hour (30 min on, 30 min off) over a time period of 1 or 2 days per site. We opted for such short recording periods with the aim of capturing the acoustic diversity of multiple sites by rotating the recorders across the landscape within the period of maximal bird vocal activity (Sugai et al., 2020). The sampling period, spanning from late April to mid-June in 2019-2021, was intended to capture not only breeding bird songs, but also the amphibian choruses at dusk and night and the beginning of acoustic activity by sound-producing insects such as orthopterans and cicadas (Grant & Samways, 2016).

In the 16 mountain sites, on the other hand, we used Song Meter Mini recorders (Wildlife Acoustics) configured to record uninterruptedly for 1 to 5 days at a sampling rate of 24 kHz. The acoustic sampling took place slightly later in the year, from mid-May to mid-July 2022, coinciding more precisely with the period of maximal vocal activity of montane birds. Finally, in the remaining 3 lowland sites we used Song Meter Micro recorders (Wildlife Acoustics) configured to record 1 minute every 15 minutes at a sampling rate of 48 kHz. The sampling took place in June and July 2022.

Among all sampling stations where acoustic data was collected, we selected the 79 for which we were able to obtain samples of the 3 times of day chosen: 0am, 7am and 1pm. The reason for choosing these three times was to maximize the total number of species detected by making sure to capture species with different patterns of daily activity. This better capture of total species richness makes it possible to test the BirdNET algorithm with a wide array of different species and allows for a more robust analysis of its general performance. The variability in recording dates, which range from late April to early July, implies that the three times of day selected can translate into slightly different moments of the day relative to sunrise and sunset. However, regardless of the specific recording date within the aforementioned range, 0am remains a fair representative of early night, 7am of early morning and 1pm of early afternoon, thus maintaining a high degree of homogeneity in the biological activity they capture across dates. Of the 237 samples analyzed, 21 were recorded in late April, 118 in May, 92 in June and 6 in early July.

***Rambouillet***

The Rambouillet area is dominated by sessile and pedunculate oaks (Quercus petraea and Q. robur) and a diverse deciduous tree community as secondary species, including common beech (Fagus sylvatica), aspen (Populus tremula), birches (Betula spp), hornbeam (Carpinus betulus), wild cherry (Prunus avium), wild service (Sorbus torminalis), ash (Fraxinus excelsior), apple (Malus sylvestris) and sweet chestnut (Castanea sativa). Conifers, most notably Scots pine (Pinus sylvestris), also cover about 20% of the forest area (Temoin, 2009).

With the aim of protecting mature forest biodiversity, a network of 141 old-growth forest (OGF) preserves equally distributed across the entire massif was designed and established in Rambouillet about 10 years ago (Temoin, 2009). These OGF preserves have been set aside from silvicultural production cycles in order to protect and support old-growth large trees, which are either left unmanaged or carefully managed with the goal of preserving biodiversity. These OGF preserves have an average area of 3 ha and include both ageing and senescent stands.

Environmental soundscapes were collected in the Rambouillet forest during the first COVID-19 lockdown in France, between April 1st and May 9th 2020. A total of 25 AudioMoth recorders were deployed in 68 forest stands (Fig. 1b). Half of these stands are located within the OGF preserve network, with 15 of them in unmanaged rewilding preserves and the other 19 in carefully-managed biodiversity-friendly preserves. The remaining 34 sites are found in the conventional production stands dominated by mature oak forests.

The acoustic sampling was conducted with a discontinuous recording setting of 30 min on / 30 min off during 3-9 consecutive days (mean = 5.8 days) and a sampling rate of 48 kHz. All AudioMoths were installed following the same procedure: a mid-size tree (oak tree, birch or hornbeam) found at >50 m from the stand edge was arbitrarily selected, then the device was attached to a low branch between 1.5 and 2 m above the ground. Once retrieved, recordings between 6:00 and 6:05, roughly corresponding to the time of maximal bird vocal activity, were selected for the acoustic analysis. More specifically, 3 such recordings (corresponding to three consecutive days) were selected for each sampling site.

***TreeBodyguards***

Our third dataset was inherited from a pan-European citizen science project born with the aim of characterizing the bottom-up and top-down trophic interactions between pedunculate oaks (Quercus robur), the caterpillars feeding on them and the natural enemies of these caterpillars (birds, mammals and arthropods) along a 19º latitudinal gradient (Valdés-Correcher et al., 2021).

Environmental soundscapes were recorded between May and July 2021 in 47 sampling sites across 17 European countries ranging from Spain to North-West Russia and covering most of the pedunculate oak geographic range (Fig. 1c). All sites are located in relatively urbanized areas meeting the single requirement of being surrounded by a wooded area of at least one hectare. The authors of the original study equipped one pedunculate oak tree per site with an Audiomoth device set up to record 30 minutes every hour (30 minutes on, 30 minutes off) at a sampling rate of 48 kHz. The automated recording started six weeks after budburst in each study area, thus synchronizing the recording period with local oak phenology, and lasted until total exhaustion of the batteries. Audiomoths were active for 9 days on average, corresponding to a total of 5920 hours of recording across all sites.

For each recording site, we selected 30 minutes of environmental soundscape recorded between 30 minutes before and 3.5 hours after sunrise. We cut the selected recordings into 10-minute subsamples and we kept the subsamples recorded on Tuesdays, Thursdays, Saturdays and Sundays. We deliberately chose to retain recordings from both working and non-working days in order to have a representative gradient of anthropogenic noise intensities. We then plotted the spectrogram of the selected subsamples using the R Seewave library in order to filter out audio samples with high levels of antropophony or geophony. Finally, we randomly selected one sample per site and day from the remaining pool of recordings, with the exception of four sites for which the four samples only covered two to three days. This makes for a total of 188 audio samples.

**Appendix S2. Manual bird identification by experts**

For the annotation of the ZA-PyGar dataset, the software Audacity was chosen because of its versatility, the convenience of its annotation export format and its overall ease of use. The annotations made with Audacity (Fig. S1) consist of 4 codes: the 6-letter alpha code for the species identified (e.g., "Tur mer" for Turdus merula, "Syl atr" for Sylvia atricapilla), the level of confidence in the identification of the species (1 for certain identifications and 0 for uncertain ones), a code for the type of sound annotated (1 for songs, 2 for static calls, 3 for flight calls, 4 for drumming, 5 for wing flapping and 6 for begging calls) and a code for the number of birds detected in the sound fragment encapsulated by the annotation (1 for 1 bird, 2 for 2 birds of the same species, 3 for more than 2 birds of the same species and 4 for 2 or more birds of different species). Distant, noisy or doubtful sounds were listened to more than once if needed. For the evaluation of BirdNET we only considered the manual identifications deemed to be certain. Identifications deemed as uncertain correspond, for the most part, to distant sounds or to sounds masked by background noise or by other birds vocalizing at the same time. Even in circumstances where a BirdNET identification coincides with an uncertain identification by the expert birder, these identifications are subject to being erroneous, so we considered it prudent not to include them in our analyses.

**Appendix S3. BirdNET configuration**

BirdNET offers the possibility to select the list of species that we want the algorithm to look for in the recording. If no species list is provided, the algorithm automatically generates one of its own by setting a minimum occurrence threshold of 0.05. This selects the list of species with an estimated 5% or higher probability of being detected in the recording based on the geographic coordinates and week of the year entered as input. In this study we used BirdNET itself to generate the list of species potentially detectable in each recording by setting a minimum occurrence threshold of 0.02, and we then feeded these lists back to the algorithm. We opted for a minimum occurrence threshold lower than the one by default because the latter filtered out a non-negligible number of species that were actually present in the recordings. We did not select a value lower than 0.02 because, even if it resulted in the additional inclusion of some of the species present in the recordings, it also ended up including a substantial number of non-present species that appreciably increased the false positive rate of the algorithm.

**Appendix S4. Clarifications about the categorization criteria used to classify BirdNET results**

The categorization criteria described in Table 2 can be interpreted as overly favorable towards BirdNET, since they imply that a correct identification of a given species takes precedence over multiple wrong identifications of the same species in the same acoustic sample. However, the most common way that BirdNET identifications are likely to be used in biodiversity monitoring projects is by generating a list of species present in a given acoustic dataset. In this context, provided that a species has been correctly detected at least once in the dataset, this species should be rightly included in the final detection list regardless of its potential misidentifications at other moments within the same dataset. Hence, we believe this categorization criterion to be an appropriate way of assessing the reliability of BirdNET as applicable to its most commonly intended use.

**Appendix S5. Clarifications about the confidence thresholds used in Figs S4-S6**

We calculated the three minimum confidence thresholds considered optimal according to the F1, F0.25 and F0.1 scores described in section 2.4. The optimal minimum confidence threshold for a given project will depend, in any case, on the importance that each user or research group assigns to minimizing FPs vs. minimizing FNs. The amount of data analyzed in this study did not allow for recommendations of optimal minimum confidence thresholds for specific species, so we only estimated the best possible values of this parameter for optimized global results when consistently used across all species. Once the optimal minimum confidence thresholds were determined for the three F-scores analyzed, we proceeded to perform species-specific precision and recall analyses for each species detected by BirdNET (in the case of precision) or identified by an expert (in the case of recall) using each of the three thresholds.

**Appendix S6. Influence of anthropophony and geophony on BirdNET performance**

For the analysis on the influence of anthropogenic and geological (e.g., wind or rainfall) sounds on BirdNET performance we used a customized version of the Normalized Difference Soundscape Index (NDSI) (Kasten et al., 2012), which provides an estimate of the preponderance of biological sounds over anthropogenic ones in a recording. According to this index, all sounds with frequencies above 2000 Hz are considered to be of biological origin, whereas sounds with frequencies between 1000 and 2000 Hz are classified as anthropogenic. It should be noted, however, that this classification criterion is fallible, since the frequencies of biological sounds partially overlap with those of anthropogenic ones, e.g., the sounds of some birds such as crows, ravens or magpies fall mostly within the range of frequencies considered to be anthropogenic. Nonetheless, despite its inability to accurately ascertain the origin of certain sounds, this index remains a fairly accurate proxy for the predominance of biological sounds over anthropogenic ones in environmental soundscapes (Bradfer-Lawrence et al., 2023). To measure the impact of both geophony and anthropophony on BirdNET performance, we customized the NDSI calculation to estimate the preponderance of biological sounds (≥2000 Hz) over all non-biological ones (<2000 Hz). This removal of the lower threshold of 1000 Hz used by default by NDSI to delimit anthropogenic sounds allowed us to capture and include a broader range of geological sounds within the non-biological category (Ross et al., 2021).

The results obtained suggest that rec_recall scores are positively correlated with the preponderance of biological sounds in relation to non-biological ones in the recordings, as measured by NDSI (Fig. S11; Spearman: rs = 0.187 and p = .004). While running in the same direction, the influence of NDSI on rec_precision and rec_F1-scores does not reach the threshold for statistical significance (Spearman, rs = 0.076 and p = 0.253 for rec_precision, rs = 0.090 and p = .173 for rec_F1-scores).

| **Category** | **ZA-Pygar** | **Rambouillet & TreeBodyguards** |
| --- | --- | --- |
| **True Positive** | Since the specific vocalization times of each species have been annotated, BirdNET identifications are only considered correct if both BirdNET and the expert identified the same species at the same time. Provided that a species has been correctly detected once in the acoustic sample, we consider the observation to be a TP even if other vocalizations of the same species were not correctly identified. | Within the acoustic sample analyzed there was a sound emitted by a bird species that the algorithm identified correctly, i.e., both BirdNET and the expert identified the same species. |
| **False Positive** | The species was mistakenly detected at least once and was not correctly detected at any moment in the acoustic sample analyzed. Sometimes a species can be correctly detected at one time and incorrectly detected at a different time within the same acoustic sample; these cases will be considered TPs. | A species was detected by BirdNET but not by the expert birder in the acoustic sample analyzed. |
| **True Negative** | A species not present in the acoustic sample was not detected by BirdNET at any time. | A species not present in the acoustic sample was not detected by BirdNET at any time. |
| **False Negative** | A species present in the acoustic sample failed to be detected by BirdNET at any moment over the sample duration. | A species present in the acoustic sample failed to be detected by BirdNET at any moment over the sample duration. |

**Table S1:** Categorization criterion followed when assessing BirdNET performance for each acoustic dataset

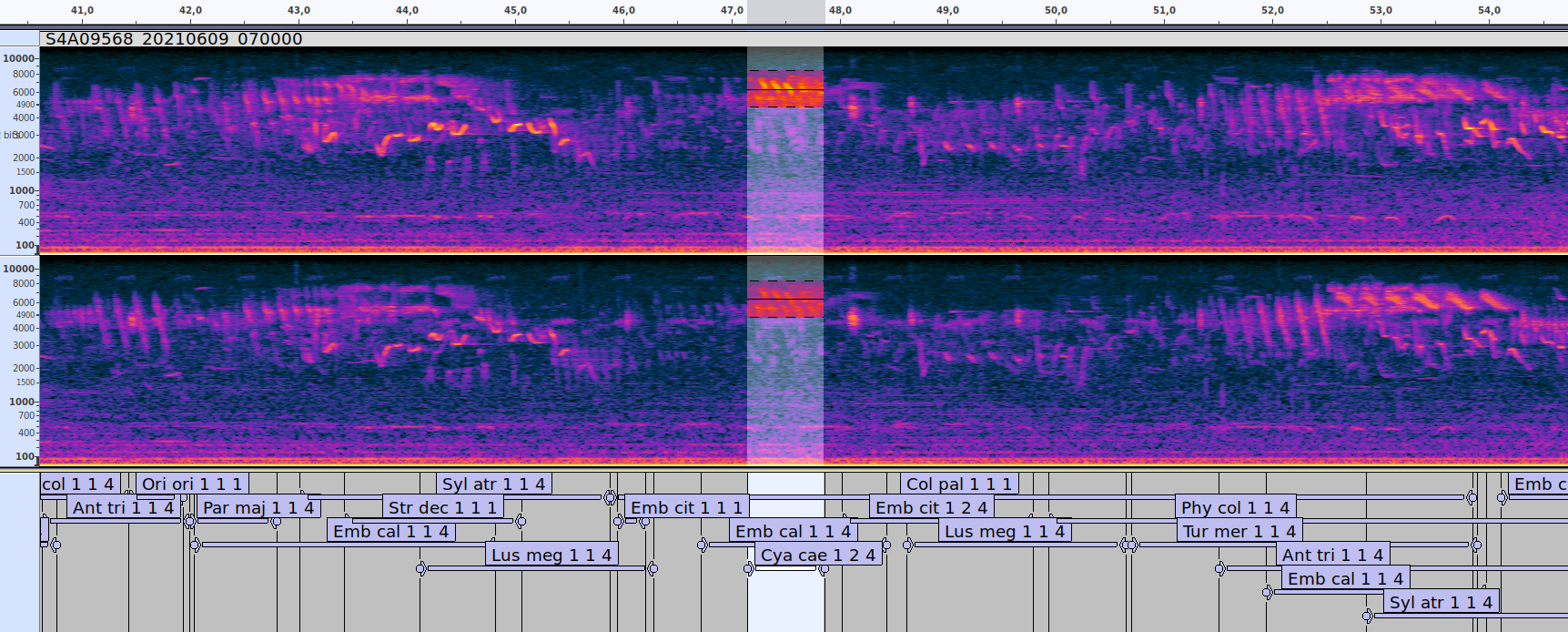

**Fig. S1:** Annotations made with Audacity: the two divisions on top show the spectrograms corresponding to the left and right channels of the recording, whereas the division at the bottom shows the manual annotations identifying all sounds present in the recording. As can be observed for the selected annotation (Cya cae 1 2 4), all annotations encapsulate a bird vocalization both in time and in frequency. Annotations consist of 4 codes: the 6-letter alpha code for the species identified (e.g. "Tur mer" for Turdus merula, "Syl atr" for Sylvia atricapilla), the level of confidence in the identification of the species (1 for certain identifications and 0 for uncertain ones), a code for the type of sound annotated (1 for songs, 2 for static calls, 3 for flight calls, 4 for drumming and 5 for wing flapping) and a code for the number of birds detected in the sound fragment encapsulated by the annotation (1 for 1 bird, 2 for 2 birds of the same species, 3 for more than 2 birds of the same species and 4 for 2 or more birds of different species).

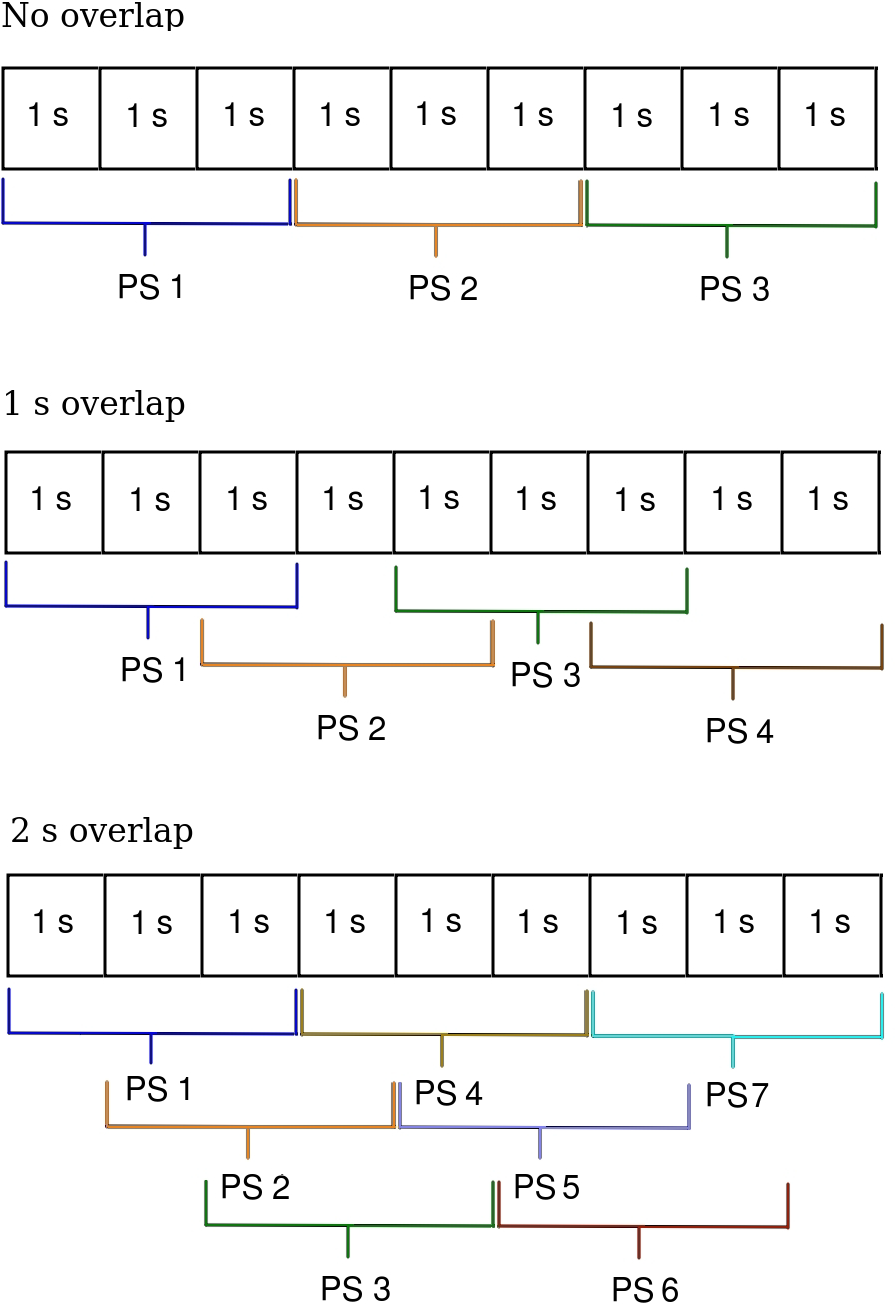
Fig. S2: Diagram illustrating the distribution of prediction segments (PS) over a given recording depending on the degree of overlap (0, 1, or 2 seconds) allowed by BirdNET.

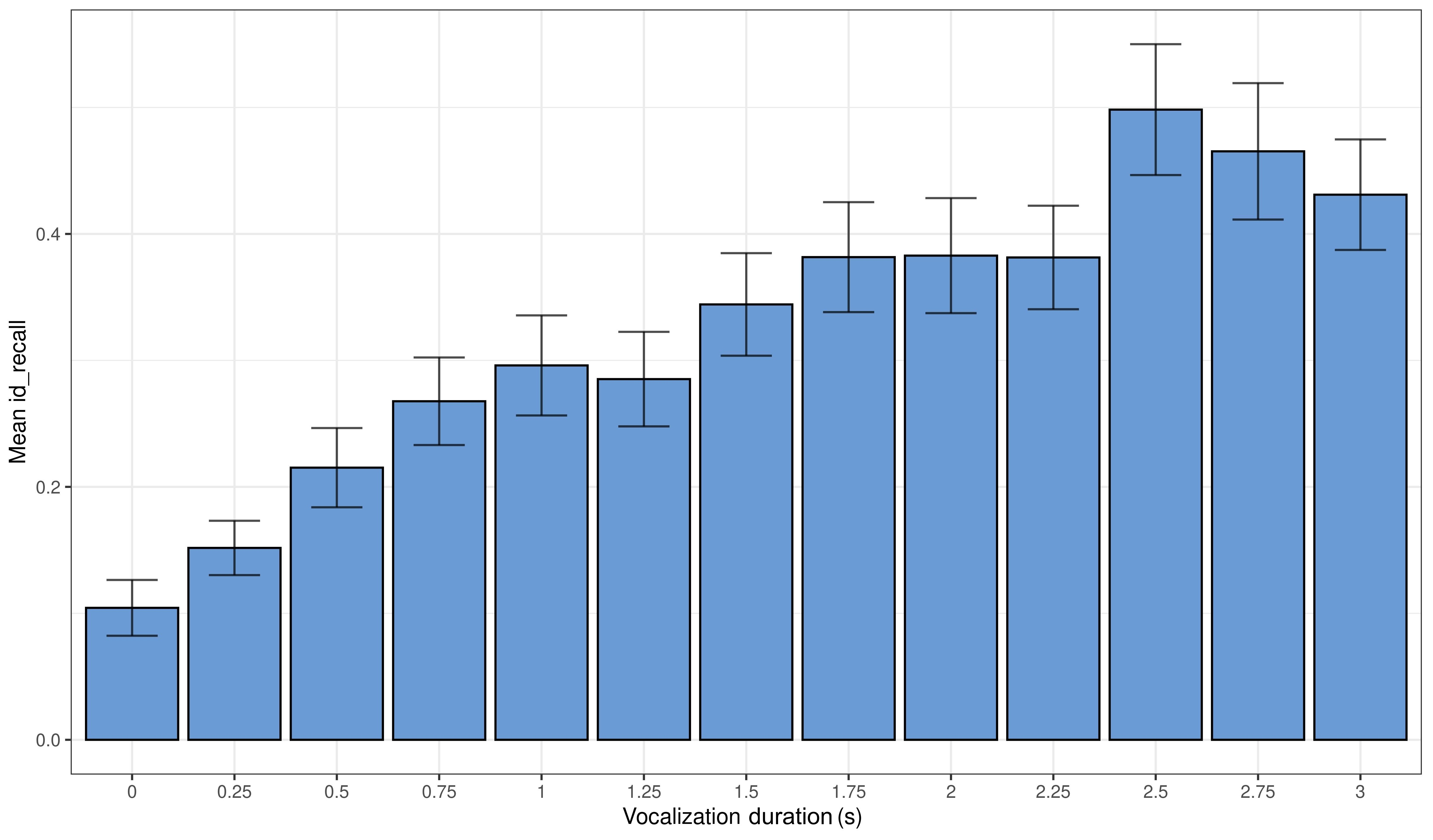

**Fig. S3:** Mean id_recall, along with the standard error bar, of BirdNET identifications for different durations of the target bird vocalization captured within the 3-second prediction segment analyzed (confidence score ≥ 0.1). Only recordings from the ZA-Pygar dataset are included.

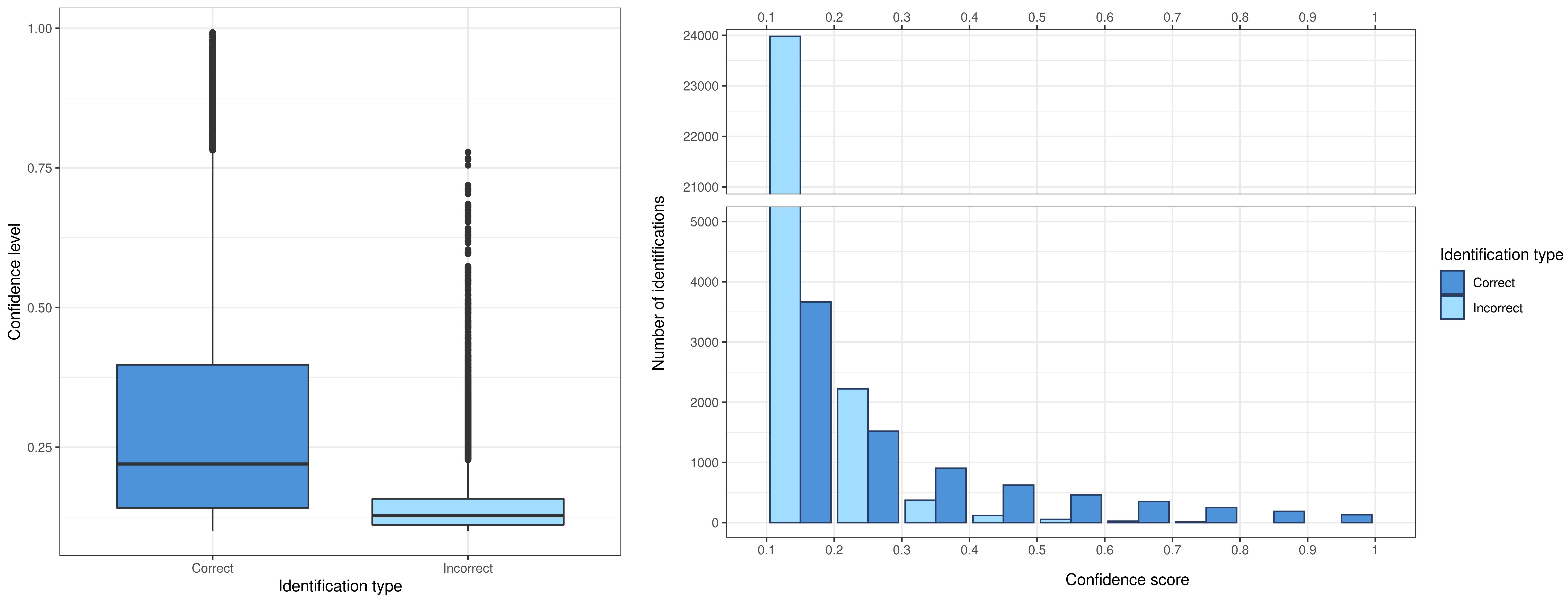

**Fig. S4:** Distribution of confidence scores in BirdNET results for both correct and incorrect identifications. Only recordings from the ZA-Pygar dataset are included.

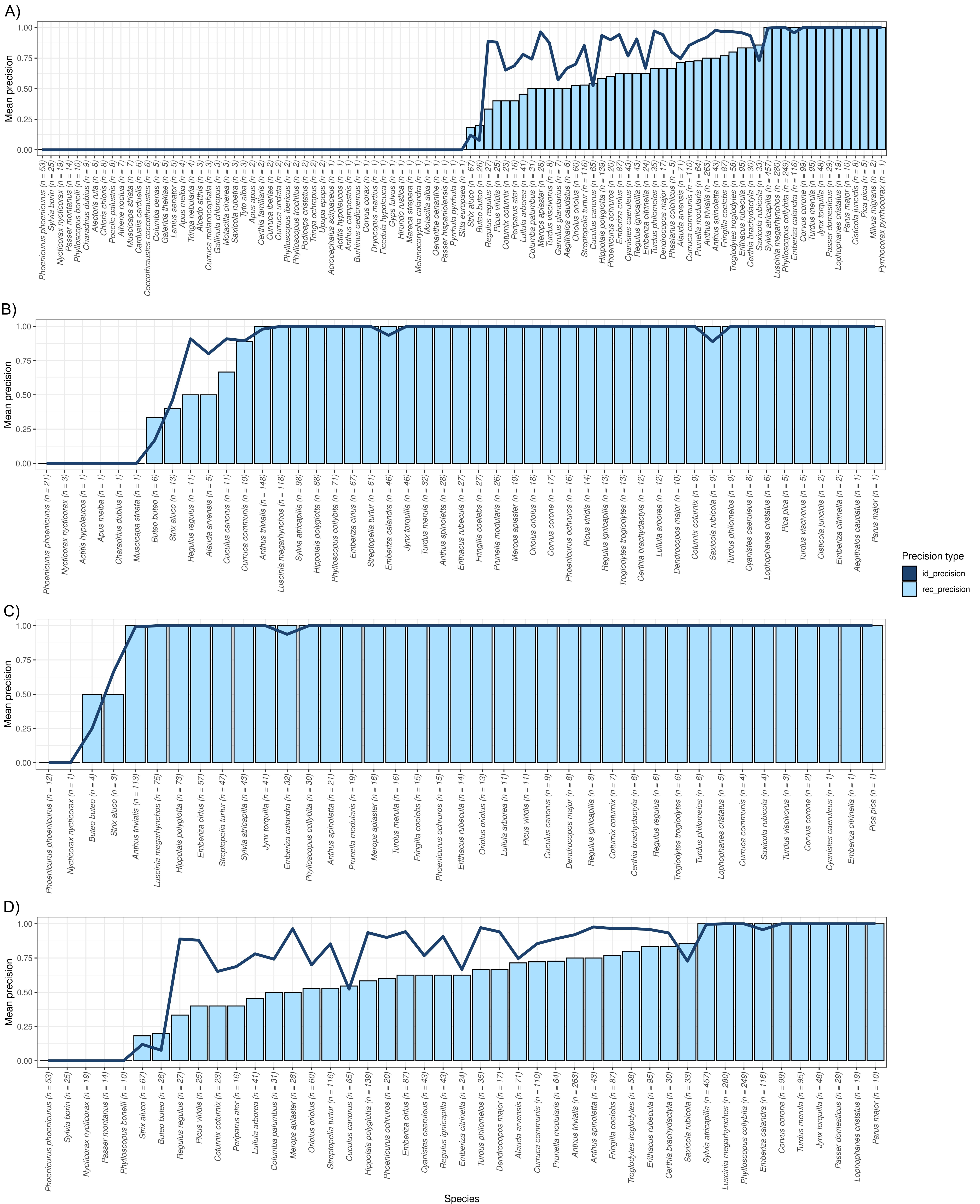

**Fig. S5:** Mean id_precision (line) and rec_precision (bars) by bird species in BirdNET identifications with confidence scores (A) ≥0.3, (B) ≥0.55, (C) ≥0.65 and (D) ≥0.3, the latter only including species with ≥10 BirdNET detections. Only recordings from the ZA-Pygar dataset are included. The number following each species name indicates the number of prediction segments where the species in question has been detected by BirdNET.

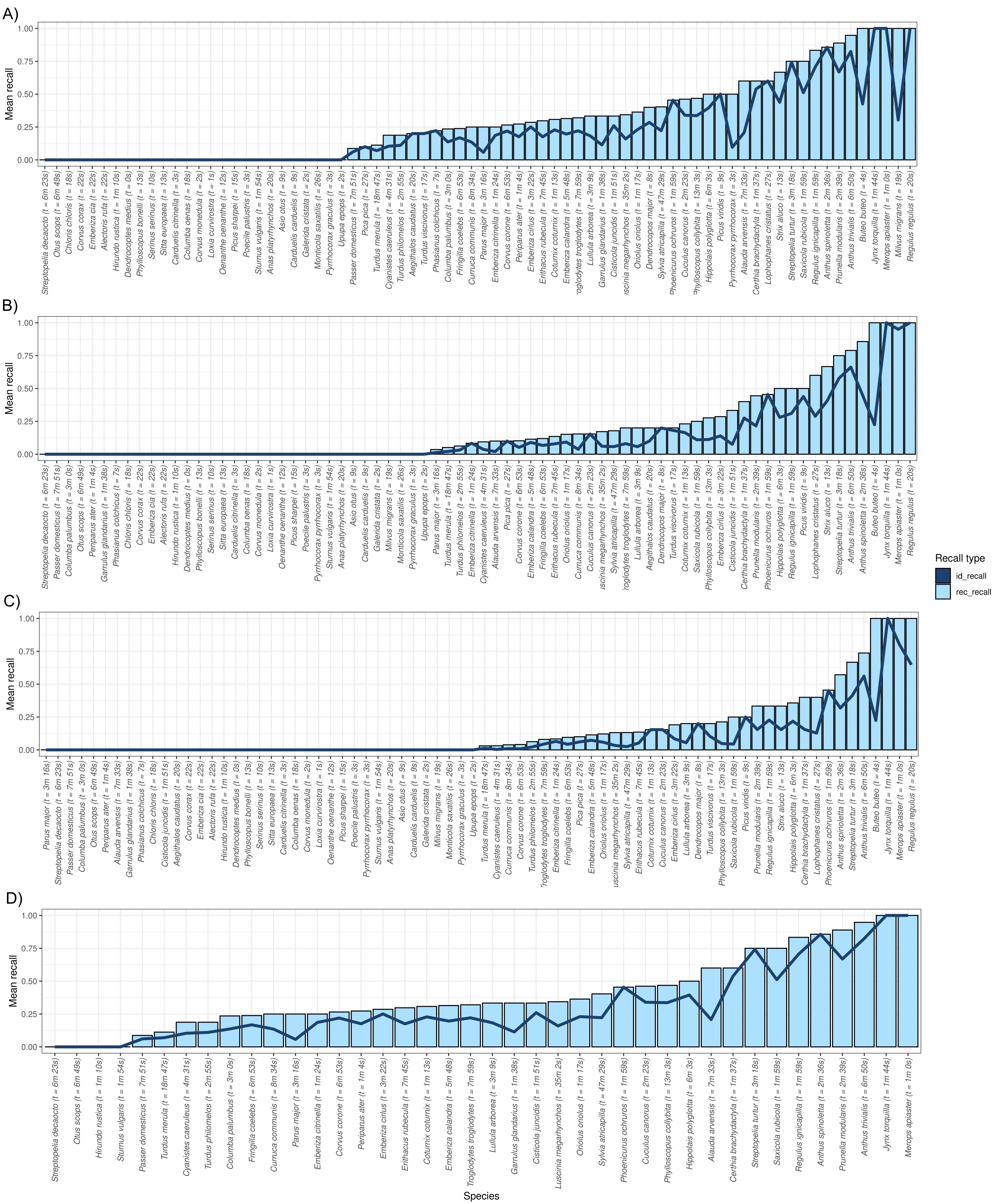

**Fig. S6:** Mean id_recall (line) and rec_recall (bars) for each bird species present in the recordings of the ZA-Pygar dataset. Only bird identifications with a confidence score (A) ≥0.3, (B) ≥0.55, (C) ≥0.65 and (D) ≥0.3 are considered, the latter plot only showing results for species with a (manually annotated) total vocalization time of ≥30 s across all recordings analyzed. Only recordings from the ZA-Pygar dataset are included.

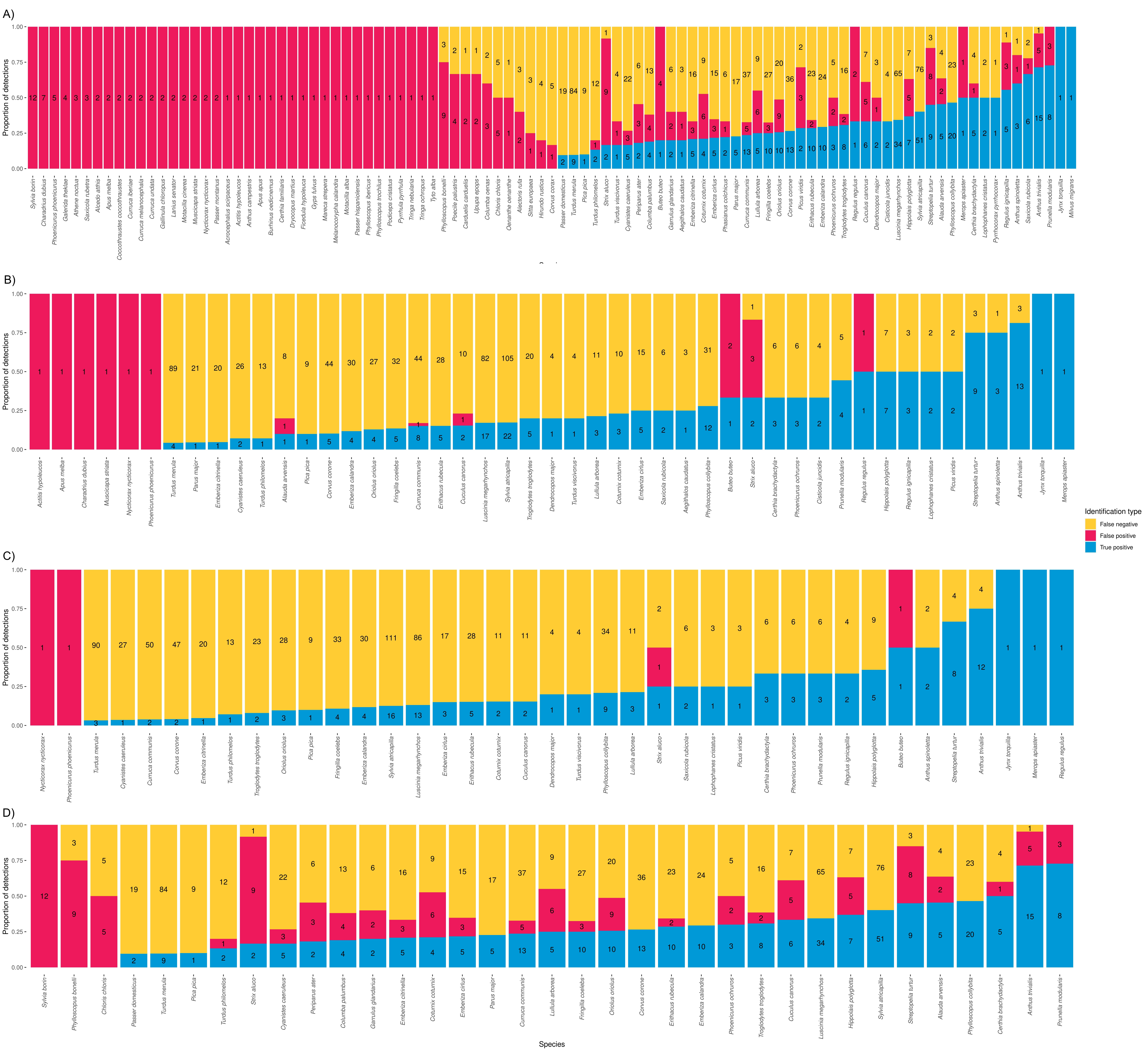

**Fig. S7:** Proportion of true positives, false positives and false negatives by bird species in BirdNET results with confidence scores (A) ≥0.3, (B) ≥0.55, (C) ≥0.65 and (D) ≥0.3, the latter only including species with ≥10 detections (either by BirdNET or by an expert). Results are calculated at the recording level and only recordings from the ZA-Pygar dataset are included.

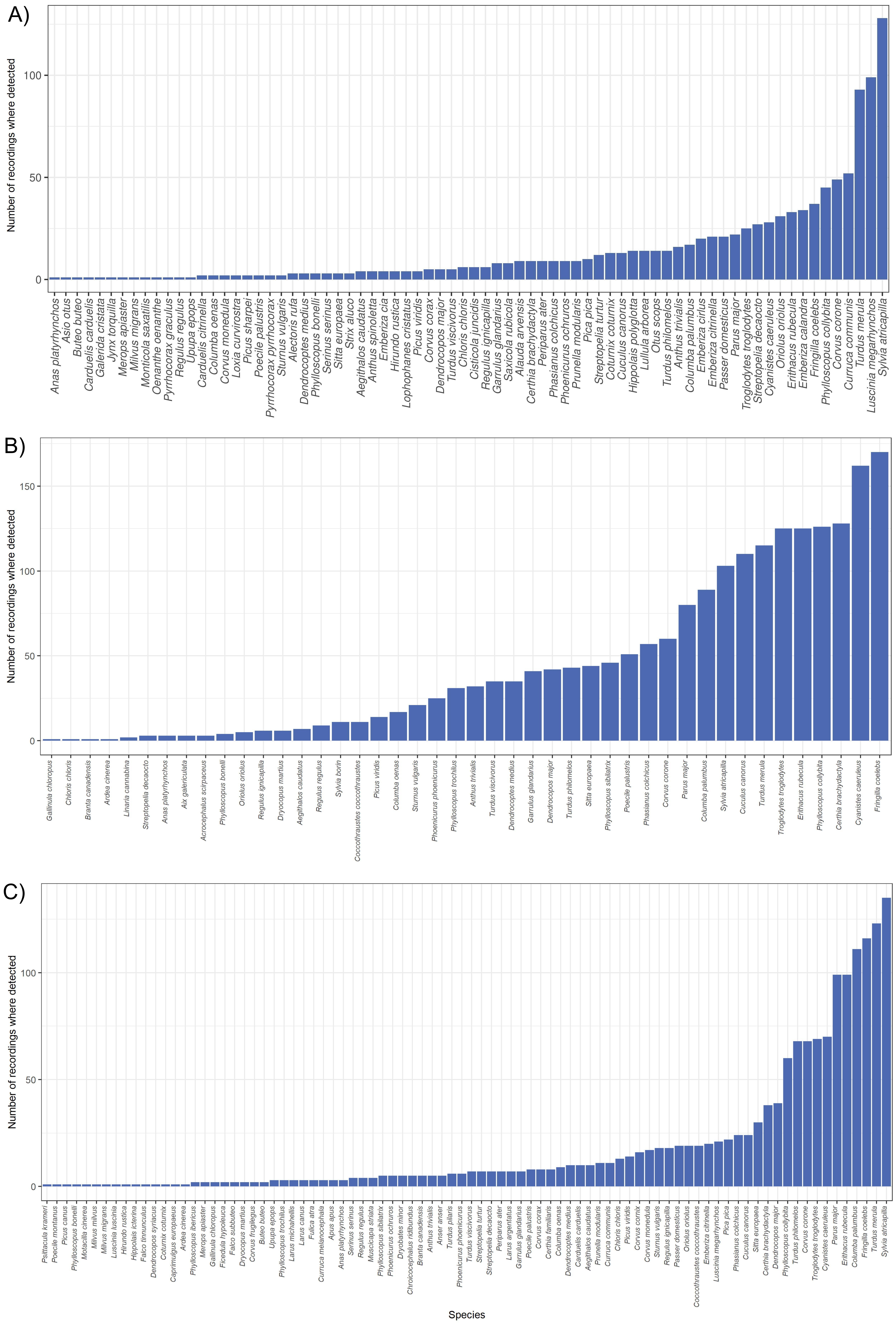

**Fig. S8:** Number of recordings where each bird species has been manually detected in the (A) ZA-Pygar, (B) Rambouillet and (C) TreeBodyguards datasets.

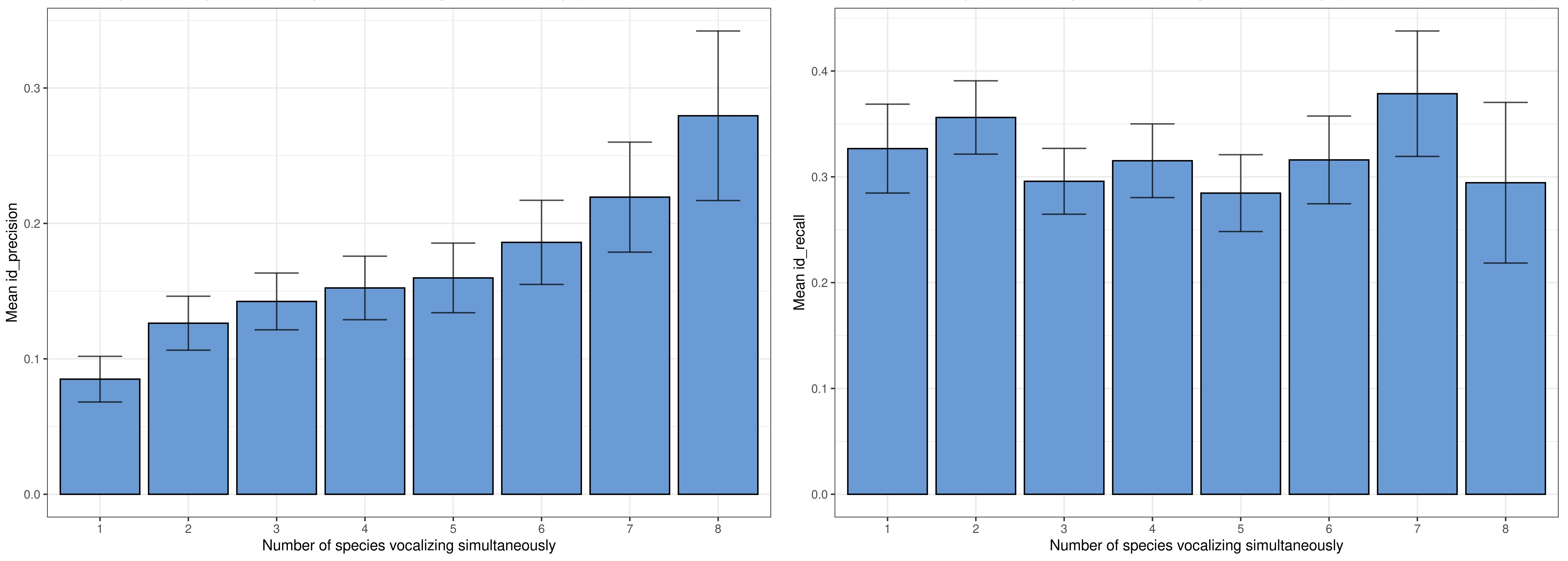
**Fig. S9:** Mean id_precision (left) and id_recall (right) scores, along with the standard error bars, depending on the number of species vocalizing simultaneously in a 3-second prediction segment (minimum confidence threshold = 0.1). Only recordings from the ZA-Pygar dataset are included.

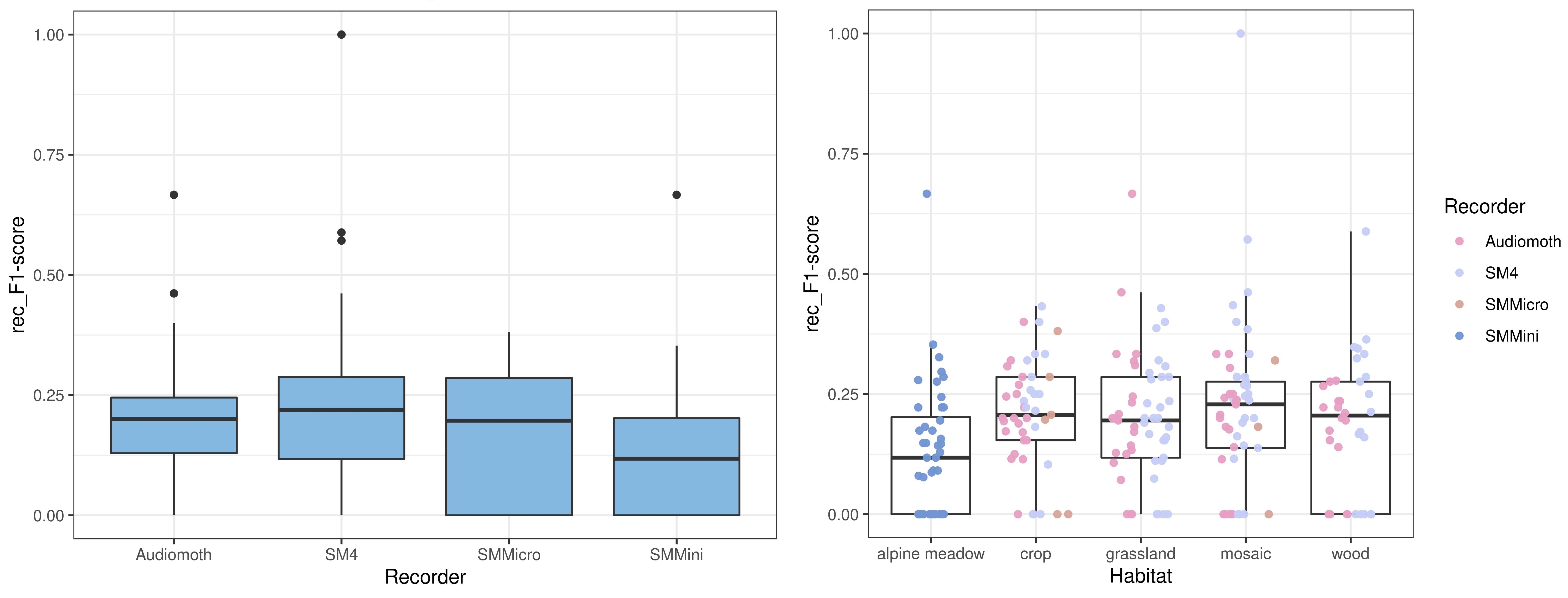

**Fig. S10:** Distribution of the rec_F1-scores obtained for each audio file analyzed (minimum confidence threshold = 0.1) depending on the recorder used (left) and the predominant habitat of the sampling site where it was recorded (right). Only recordings from the ZA-Pygar dataset are included.

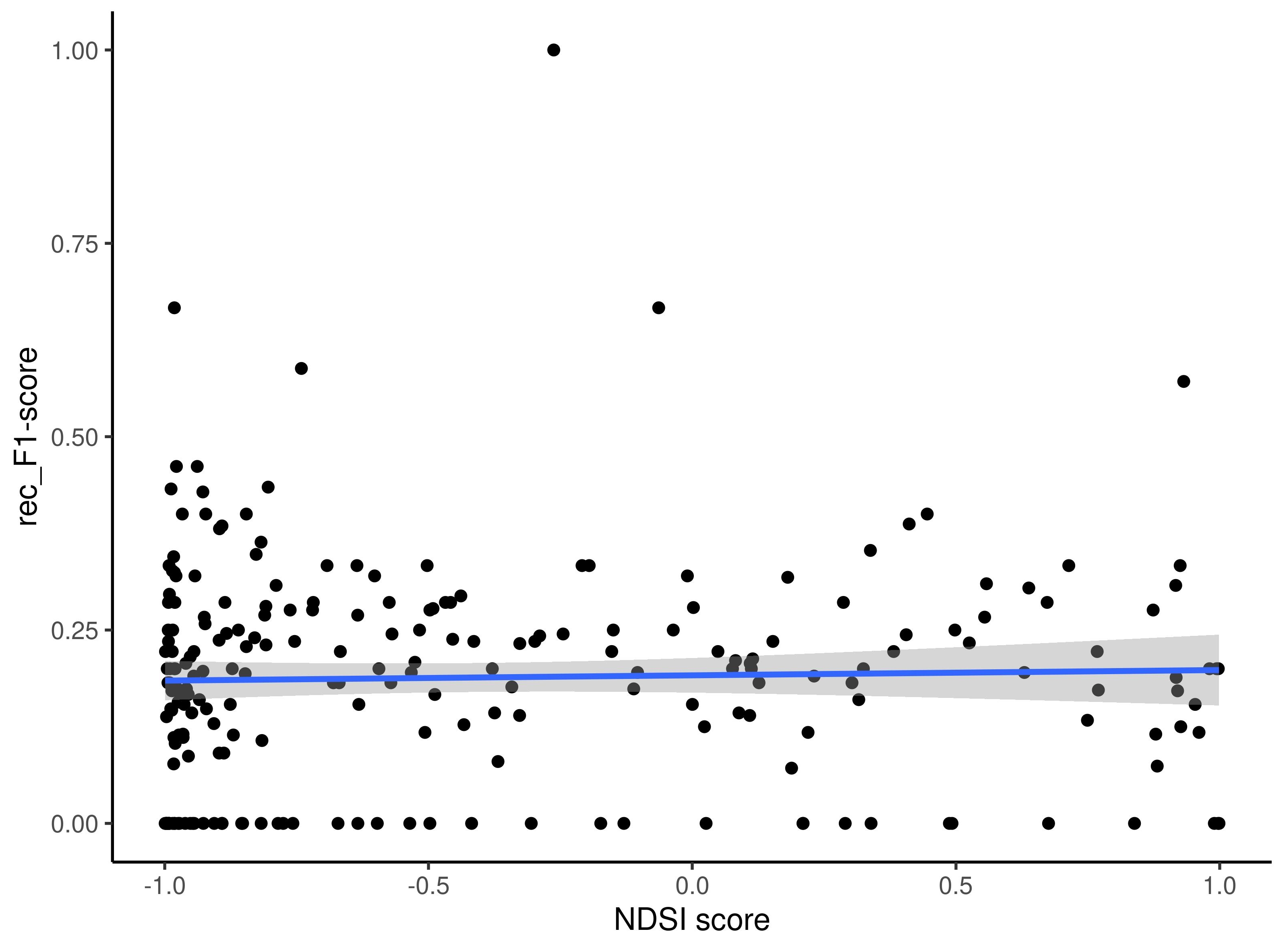

**Fig. S11:** Relationship between BirdNET performance and the predominance of biological sounds over non-biological ones for every recording. More specifically, the mean rec_precision (top-left), rec_recall (top-right) and rec_F1-scores (bottom) obtained for each recording (minimum confidence threshold = 0.1) are plotted against the corresponding NDSI scores. Only recordings from the ZA-Pygar dataset are included.
